# Predicting Brain Volumes from Anthropometric and Demographic Features: Insights from UK Biobank Neuroimaging Data

**DOI:** 10.1101/2025.06.22.660902

**Authors:** Kimia Nazarzadeh, Simon B. Eickhoff, Georgios Antonopoulos, Lukas Hensel, Caroline Tscherpel, Vera Komeyer, Federico Raimondo, Christian Grefkes, Kaustubh R. Patil

**Affiliations:** Department of Neurology, University Hospital Cologne, University of Cologne, Cologne, Germany; Institute of Neuroscience and Medicine, Brain & Behaviour (INM-7), Research Centre Jülich, Jülich, Germany; Institute of Neuroscience and Medicine, Cognitive Neuroscience (INM-3), Research Centre Jülich, Jülich, Germany; Institute of Systems Neuroscience, Medical Faculty and University Hospital Düsseldorf, Heinrich Heine University, Düsseldorf, Germany; Goethe University Frankfurt, Department of Neurology, Frankfurt University Hospital, Frankfurt am Main, Germany; Department of Biology, Faculty of Mathematics and Natural Sciences, Heinrich Heine University Düsseldorf, Düsseldorf, Germany

**Keywords:** brain volume, sexual dimorphism, machine learning, UK Biobank, structural MRI, anthropometrics

## Abstract

Brain size measures are well-studied and often treated as a confound in volumetric neuroimaging analyses. Yet their relationship with body anthropometric measures and demographics remains underexplored. In this study, we examined those relationships alongside age- and sex-related differences in global brain volumes. Using brain magnetic resonance imaging (MRI) of healthy participants in the UK Biobank, we derived global measures of brain morphometry, including total intracranial volume (TIV), total brain volume (TBV), gray matter volume (GMV), white matter volume (WMV), and cerebrospinal fluid (CSF). We extracted these measures using the Computational Anatomy Toolbox (CAT) and FreeSurfer. Our analyses were structured in three approaches: across-sex analysis, sex-specific analysis, and impact of age analysis. Employing machine learning (ML), we found that TIV was strongly predicted by sex (across-sex *r* = 0.68), reflecting sexual dimorphism. On the other hand, TBV, GMV, WMV, and CSF were more sensitive to age, with higher prediction accuracy when age was included as a feature, highlighting age-related changes in the brain structure, such as fluid expansion. Sex-specific models showed reduced TIV prediction (*r* ≈ 0.25) but improved TBV accuracy (*r* ≈ 0.44), underscoring sex-specific body-brain relationships. Anthropometrics enhanced prediction but only subsidiary to age and sex. These findings advance our understanding of brain-body scaling relationships and underscore the necessity of accounting for age and sex in neuroimaging studies of brain morphology.

## 1. Introduction

The intricate interplay between brain morphology and body characteristics represents a fundamental aspect of neuroimaging research, revealing how brain structure varies with respect to factors such as anthropometrics, age, sex, genetics, and environmental conditions (Gurholt et al. 2021). Understanding these complex body-brain relationships is critical for advancing our knowledge of brain structure and function and for improving the precision and utility of neuroimaging studies, particularly in the context of health, aging, and disease (Gurholt et al. 2021). Key global volumetric measures of brain morphology, including total intracranial volume (TIV), representing the total cranial capacity; total brain volume (TBV), the sum of whole-brain gray matter volume (GMV) and whole-brain white matter volume (WMV), and cerebrospinal fluid volume (CSF) are critical for understanding how brain structure varies with individual factors such as age, sex, and body characteristics (Raz and Rodrigue 2006; Bayat et al. 2012). Multiple studies have demonstrated that these brain volumes are associated with anthropometric features such as height, weight, and body mass index (BMI), waist circumference (WC), and hip circumference (HC), with varying effects across sexes and age groups. For instance, height and weight are positively associated with brain volumes, suggesting that larger body sizes correspond to larger brain volumes. Conversely, higher WC, an indicator of central obesity, is associated with reduced TBV and GMV, particularly in frontal, temporal, and limbic regions (Ward et al. 2005; Bayat et al. 2012). Similarly, higher HC, which reflects lower-body fat distribution, has been associated with larger brain volumes, although the underlying mechanisms and implications of this relationship have not been fully understood (Eboh et al. 2016).

Age, as a ubiquitous biological process, is a pivotal factor shaping brain volumes across the lifespan, driving significant structural changes, making it a primary risk factor for various brain disorders (Smith et al. 2007). These age-related alterations, modulated by other factors such as sex, genetics, and environmental factors, manifest distinctly across brain compartments. Evidence reveals distinct patterns of change for each brain volume (Smith et al. 2007; Chen et al. 2007; Ruigrok et al. 2014). TIV increases during childhood, then stabilizes in adulthood due to the fixed cranial structure. In contrast, TBV and its components decline markedly with aging (Mills et al. 2016). GMV decreases linearly from early adulthood, with studies showing pronounced reductions in the frontal cortex by middle age and loss across multiple regions over time. WMV follows a different trajectory, initially increasing into early adulthood before declining later in life, with evidence indicating deterioration beginning in young adulthood and progressing across various brain areas. Meanwhile, CSF volume increases with age, a consequence of brain tissue atrophy. These widespread and progressive age-related changes emphasize the dynamic and differential effects of aging on brain structure. Sex, further, as a fundamental biological factor, significantly influences patterns of brain structure and aging. Sexual dimorphism in brain morphology is well-established, with males generally exhibiting larger absolute global brain volumes than females. On average, males have approximately greater TIV (12%), TBV (11%), GMV (9%), WMV (13%), and CSF (11.5%) compared to females (Chen et al. 2007; Ruigrok et al. 2014; Eliot et al. 2021). These percentages reflect larger brain volumes in males, which generally align with greater body size during development, as larger bodies typically correspond to proportionally larger brains. However, when controlling for overall body size, these differences diminish significantly (Ruigrok et al. 2014; Mills et al. 2016; Eliot et al. 2021). Regionally, females tend to have a larger fronto-parietal cortex, while males show a larger occipito-temporal cortex (Ritchie et al. 2018). In contrast, findings for subcortical structures, such as the hippocampus, remain inconsistent across studies (Chen et al. 2007; Ruigrok et al. 2014; Lotze et al. 2019). Understanding such sex-specific differences in global brain volumes is essential for accurately interpreting neuroimaging data and could inform clinical practices regarding neurodegenerative diseases that exhibit sex-specific patterns of progression (Ferretti et al. 2018).

Despite these insights, a critical gap remains regarding which anthropometric measures, alongside age and sex, are predictive of brain volumes at the individual level. Addressing this gap is vital for mapping the normative body-brain relationships and identifying potential biomarkers of brain health and aging, which could guide early detection and intervention strategies for neurodegenerative conditions with sex- and age-specific profiles. Motivated by this need, our study employs Machine learning (ML) to predict brain volumes derived from structural magnetic resonance imaging (MRI) data using anthropometric and demographic features in a large cohort from the UK Biobank (*N* = 21,807). We asked two key questions: to what degree can anthropometric features predict brain volumes compared to the dominant influences of age and sex, and how do age and sex modulate the predictive power of anthropometrics across different brain volumes? These questions matter because unraveling the relative contributions of these factors could refine our understanding of brain allometry—the proportional scaling of brain and body—and reveal association of demographic and physical traits with brain structure across the lifespan. We hypothesize that TIV will be more strongly predicted by sex due to dimorphism in body proportions, while TBV, GMV, WMV, and CSF are expected to exhibit greater sensitivity to age, reflecting their susceptibility to age-related atrophy. The potential contribution of anthropometric measures to the prediction of these brain volumes will also be explored. By applying ML to assess prediction accuracies in age- and sex-related analyses, we seek to provide a comprehensive framework that clarifies these relationships and sets the groundwork for future studies linking brain morphology to health outcomes, such as cognitive decline or disease susceptibility.

## 2. Material and methods

### 2.1. Participants and data

Data for this study were obtained from the UK Biobank, a comprehensive database with over 500,000 adult participants recruited from 22 assessment centers across the United Kingdom (UK). The baseline assessment took place between 2006 and 2010. In 2014, a subset of participants was invited to undergo brain imaging. This imaging assessment included a wide range of demographic and anthropometric measurements. The healthy individuals were defined based on inclusion and exclusion criteria consistent with the International Classification of Diseases, 10th revision (ICD-10), and the detailed exclusion criteria listed in Table S1. To ensure the completeness of the dataset, only participants with complete data for demographics, anthropometrics, and brain imaging were included in the final sample (*N* = 21,807; females *N* = 11,576, males *N* = 10,231).

### 2.2. Demographic and anthropometric assessments

Sex was determined based on self-reported information (Data-Field 31). Age was calculated as the difference between the participant’s birth and assessment dates. Anthropometric measurements were collected during the physical measures phase of the assessment. These included weight (Data-Field 21002), WC (Data-Field 48), HC (Data-Field 49), seated height (Data-Field 51), seating box height (Data-Field 3077), and standing height (Data-Field 50).

### 2.3. Brain Volume

We calculated absolute brain volumes using T1-weighted (T1w) MRI data from the UK Biobank first imaging assessment visit, acquired with 3T scanners following the UK Biobank Imaging Protocol (UK Biobank Imaging Protocols). T1w images, initially preprocessed with quality control by the UK Biobank (Alfaro-Almagro et al. 2018), were retrieved and converted into a DataLad dataset for provenance tracking (Halchenko et al. 2021). For primary volumetric analysis, we used the Computational Anatomy Toolbox (CAT) version 12.7 (Gaser et al. 2024). Supplementary analyses were conducted using brain volumes derived from FreeSurfer ASEG segmentation, provided by the UK Biobank. We compared brain volumes derived from CAT and FreeSurfer to investigate how these distinct preprocessing approaches might influence brain volume estimates and subsequent predictive modeling outcomes. CAT employs a voxel-based morphometry framework with an explicit head model, which may improve spatial normalization accuracy and sensitivity to subtle structural variations, potentially benefiting models reliant on precise anatomical alignment. In contrast, FreeSurfer uses a surface-based approach with detailed tissue segmentation, which could enhance volume accuracy for tissue-specific predictions. By examining these differences, we aimed to assess their impact on the reliability and generalizability of our predictive models. The following absolute volumes were analyzed from FreeSurfer-processed UK Biobank fields: total intracranial volume (TIV, Data-Field 26521), total brain volume (TBV, Data-Field 25010), gray matter volume (GMV, Data-Field 25006), white matter volume (WMV, Data-Field 25008), and cerebrospinal fluid volume (CSF, Data-Field 26527).

## 3. ML analysis

### 3.1. Feature configurations

In this study, we examined the relationships between brain morphology (i.e., TIV, TBV, GMV, WMV, CSF) and various anthropometric measurements, including weight, WC, HC, seated height, and seating box height, and demographic factors such as sex and age. We utilized a factorial design, incorporating two primary independent factors, sex and age, structured across multiple analytical dimensions. The sex factor was evaluated in two distinct configurations: an across-sex analysis, where features of males and females were combined and sex was included as an additional feature, and a within-sex analysis, where data were analyzed separately for each sex. This dual approach enabled us to assess both the independent and interactive effects of anthropometric features, age, and sex on brain volumes from both CAT and FreeSurfer pipelines (see Section 2.3). For the age factor, we employed three analytical strategies to examine its impact: (1) including age as a feature, (2) excluding age from the feature list, and (3) removing age as a confound. These strategies were applied consistently in both across-sex and within-sex analyses. The across-sex analysis facilitated the exploration of age- and sex-related effects on brain morphology in the entire study population, while the within-sex analysis allowed us to identify sex-specific patterns and investigate how age modulates the relationships between anthropometric measurements and brain volumes within each sex group. By integrating these factorial dimensions, we aimed to provide a comprehensive evaluation of how anthropometric features, age, and sex interact to shape brain morphology in our study cohort.

### 3.2. Model training and performance evaluation

To predict brain volumes from anthropometric and demographic features, the data from both CAT and FreeSurfer preprocessing pipelines (see Section 2.3) were split into training (90%) and test (10%) datasets for both across-sex and within-sex analyses (Table 1). We employed linear support vector machine (SVM) and random forest (RF) regression models to predict brain volumes using feature combinations of anthropometric measurements (weight, WC, HC, seated height, and seating box height), age, and sex as structured by the factorial design detailed in Section 3.1. Model performance was evaluated using Pearson’s correlation coefficient (*r*), the coefficient of determination (*R*^2^), and mean absolute error (*MAE*). To ensure robust generalization estimates, we implemented 10 times repeated 10-fold (10×10-fold) cross-validation (CV) using the Julearn machine learning library (version 0.2.7; https://juaml.github.io/julearn/) (Hamdan et al. 2023), which builds on the scikit-learn library (sklearn) (Pedregosa et al.). For the linear SVM, the hyperparameter *C* was determined heuristically as 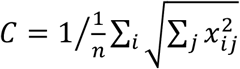 where *n* represents the number of subjects. As a nonlinear method, we also trained an RF regression model using the sklearn (version 1.2.1), with 100 trees, a minimum of 2 samples per split, the square root (sqrt) of the total number of features as the maximum number of features considered for the best split and bootstrapping of the training samples (true) as the hyperparameters (default settings in this version of sklearn). We compared the performance of the linear SVM and RF models from the 10×10-fold CV procedure using the training dataset. The model exhibiting superior performance in CV was selected for further analysis. Feature importance (FI) scores were then derived from the final models trained on the full training dataset (90% of the data) to quantify the contribution of individual features to the model outputs. For the linear SVM, FI scores were based on the coefficient parameter (scikit-learn model’s “.coef_”). Finally, we used the 10% hold-out test set to predict brain volumes and compared the predicted values to the true values, confirming the model’s performance on unseen data beyond the CV framework.

**Table 1.** Summary of the imaging population characteristics.

| Dataset | Train dataset |  |  | Test dataset |  |  |
| --- | --- | --- | --- | --- | --- | --- |
|  | Both sexes | Female | Male | Both sexes | Female | Male |
| Number | 19,625 | 10,418 | 9,207 | 2,182 | 1,158 | 1,024 |
| Age, mean (SD) | 63.32 (7.5) | 62.66 (7.28) | 64.07 (7.68) | 63.27 (7.44) | 62.29 (7.22) | 64.37 (7.54) |
| Weight, mean (SD) | 75.11 (14.74) | 68.20 (12.53) | 82.93 (13.06) | 74.84 (14.6) | 68.13 (12.48) | 82.43 (13.01) |
| WC, mean (SD) | 87.18 (12.36) | 81.72 (11.39) | 93.35 (10.35) | 86.97 (12.24) | 81.63 (11.18) | 93.01 (10.45) |
| HC, mean (SD) | 100.32 (8.52) | 100.25 (9.56) | 100.39 (7.16) | 100.19 (8.37) | 100.22 (9.26) | 100.16 (7.23) |
| Seated height, mean (SD) | 131.47 (7.26) | 127.33 (4.86) | 136.16 (6.65) | 131.54 (7.33) | 127.46 (5.05) | 136.15 (6.76) |
| Box height, mean (SD) | 42.11 (4.23) | 40.81 (3.09) | 43.59 (4.82) | 42.16 (4.34) | 40.77 (3.22) | 43.73 (4.88) |

## 4. Results and Discussion

In this manuscript, the results and discussion are presented in a combined section to facilitate a more direct interpretation of the findings. This integrated format allows each result to be immediately contextualized and discussed, improving clarity and avoiding redundancy.

### 4.1. Association between age and body-brain structure

Correlation analyses between age, anthropometric measurements, and brain volumes in female and male cohorts revealed complex age-related relationships in body-brain associations (Figure 1). WC (Figure 1B) showed weak positive correlations with age (females: *r* = 0.06, *p* < 0.001; males: *r* = 0.04, *p* = 0.0003), suggesting a subtle increase in visceral fat distribution as individuals age. The statistical significance of these correlations with small effect sizes likely reflects the large sample size rather than a meaningful biological association. Despite their weakness, these trends align with prior research documented a tendency for abdominal fat accumulation with advancing age, attributed to metabolic and hormonal changes (Kuk et al. 2009; Frank et al. 2019; Ponti et al. 2020). Caution is warranted in interpreting these weak associations, and further research may clarify their relevance. In contrast, weight, hip circumference, seated height, and standing height (Figure 1A and C-E) exhibited negative correlations. Among these, seated height (females: *r* = −0.25, *p* < 0.001; males: *r* = −0.23, *p* < 0.001) and standing height (females: *r* = −0.20, *p* < 0.001; males: *r* = −0.20, *p* < 0.001) exhibited the strongest declines, consistent with established patterns of height loss due to vertebral compression, bone mass reduction, and an excessive forward rounding of the upper back (kyphosis) (Cummings and Melton 2002; Volpi et al. 2004). On the other hand, weight demonstrated slightly lower negative correlations (females: *r* = −0.10, *p* < 0.001; males: *r* = −0.16, *p* < 0.001), likely reflecting the progressive loss of muscle mass (sarcopenia) that occurs with aging, as muscle, being denser than fat, contributes significantly to body weight (Volpi et al. 2004; Kuk et al. 2009). HC showed a much weaker negative correlations (females: *r* = −0.03, *p* = 0.0026; males: *r* = −0.06, *p* < 0.001) suggesting that fat redistribution in the hips occurs more slower than in the abdomen, potentially preserving hip shape in aging relative to some body regions with more pronounced fat accumulation (Kuk et al. 2009). These findings illustrate a complex interplay of anthropometric changes that may influence age-related alterations, highlights the importance of considering a range of body measurements when assessing the impact of aging on brain morphology.

**Figure 1.**
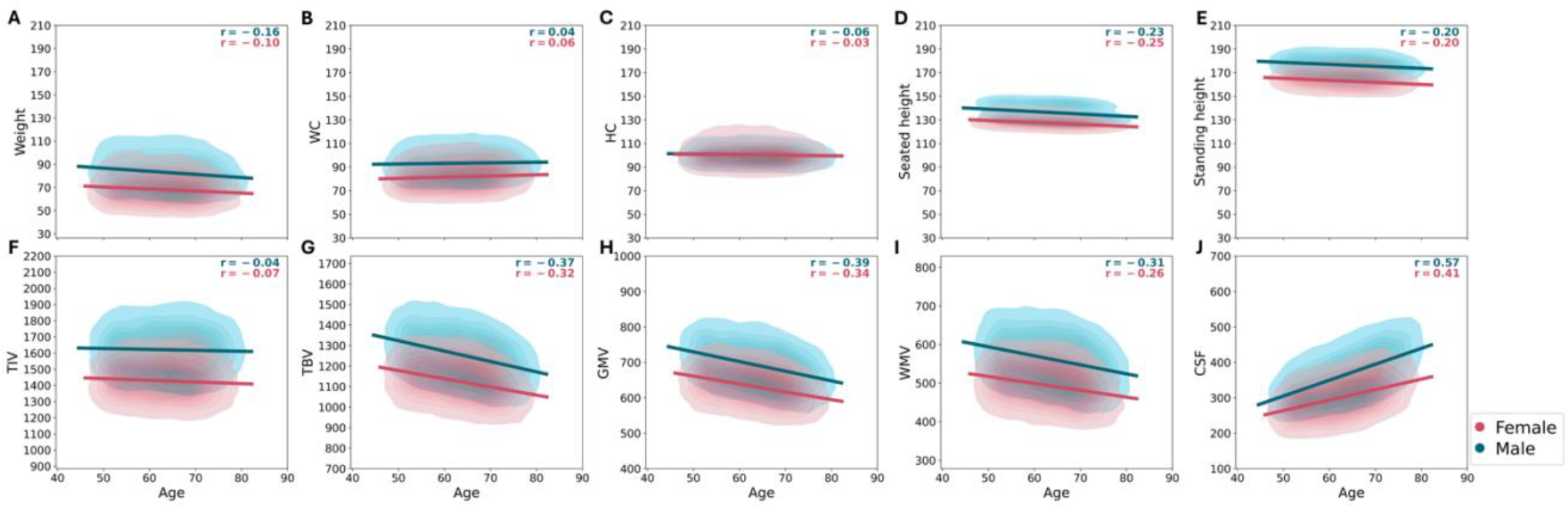
Association between age and anthropometric measurements and brain volumes, separated by sex. Panels (A) to (E) display the relationships between age and anthropometric measurements: (A) weight, (B) waist circumference (WC), (C) hip circumference (HC), (D) seated height, and (E) standing height. Panels (F) to (J) show the correlations between age and brain volumes: (F) total intracranial volume (TIV), (G) total brain volume (TBV), (H) gray matter volume (GMV), (I) white matter volume (WMV), and (J) cerebrospinal fluid volume (CSF).

Brain volumes displayed marked age-related trajectories (Figure 1F-J). TIV (Figure 1F) showed a small negative correlation with age (females: *r* = −0.07, *p* < 0.001; males: *r* = −0.04, *p* < 0.001), indicating that overall cranial capacity remains relatively stable across the lifespan. This stability is expected, as TIV includes bone structure, which is developed during childhood and remains constant throughout adulthood despite age-related changes in brain volumes (Smith et al. 2007). In contrast, TBV (Figure 1G), GMV (Figure 1H), and WMV (Figure 1I) demonstrated more pronounced decline with age. GMV exhibited the strongest negative correlations (females: *r* = −0.34, *p* < 0.001; males: *r* = −0.39, *p* < 0.001), suggesting a decline likely due to neuronal loss and synaptic pruning associated with normal aging (Gennatas et al. 2017). WMV displayed moderate negative correlations (females: *r* = −0.26, *p* < 0.001; males: *r* = −0.31, *p* < 0.001), less pronounced than GMV but still significant, potentially impacting neural communication efficiency in older adults. TBV, the sum of GMV and WMV, mirrored the declines in these volumes. Conversely, CSF (Figure 1J) increased sharply (females: *r* = 0.41, *p* < 0.001; males: *r* = 0.57, *p* < 0.001), more prominently in males, reflecting fluid expansion as brain tissue atrophies (Yamada et al. 2023). This increase may support intracranial pressure maintenance as brain tissue volumes shrink.

### 4.2. Brain volume predictive modelling

The primay analysis utilized CAT-derived data with the linear SVM model to predict brain volumes using anthropometric measurments and age. This approach was applied to both across-sex and within-sex analyses, with age included as a feature in the model (see Section 3.1). The linear SVM demonstrated robust predictive performance across all brain volumes outperforming the RF model yielding higher Pearson’s r, R^2^, and low MAE in 10×10-fold CV (Table 2). Supplementary analyses extended these findings by applying both linear SVM and RF models to FreeSurfer-derived volumes, again maintaining age as a feature (see Supplementary Tables S2 and S3).

**Table 2.** The performance of linear SVM models on the training dataset for brain volumes.

| Data | Brain volume | Scores (median) | Linear SVM |  |  |
| --- | --- | --- | --- | --- | --- |
|  |  |  | Both sexes | Female | Male |
| CAT | TIV | Pearson's $r$ | 0.67 | 0.24 | 0.24 |
| | | $R^2$ | 0.44 | 0.06 | 0.06 |
|  |  | MAE | 87.98 | 82.24 | 94.81 |
| | TBV | Pearson's $r$ | 0.64 | 0.4 | 0.43 |
| | | $R^2$ | 0.41 | 0.16 | 0.19 |
|  |  | MAE | 70.71 | 66.03 | 75.52 |
| | GMV | Pearson's $r$ | 0.62 | 0.41 | 0.45 |
| | | $R^2$ | 0.39 | 0.17 | 0.2 |
|  |  | MAE | 36.41 | 34.33 | 38.6 |
| | WMV | Pearson's $r$ | 0.6 | 0.33 | 0.37 |
| | | $R^2$ | 0.36 | 0.11 | 0.13 |
|  |  | MAE | 40.15 | 37.31 | 43.24 |
| | CSF | Pearson's $r$ | 0.67 | 0.43 | 0.57 |
| | | $R^2$ | 0.45 | 0.18 | 0.33 |
|  |  | MAE | 38.24 | 36.78 | 39.22 |

#### 4.2.1. Prediction of head size (TIV)

The prediction of TIV using anthropometric measurements, age, and sex revealed intricate relationships shaped by biological and developmental factors. Using CAT-derived data with the linear SVM model, including age as a feature, we analyzed both across-sex and within-sex cohorts (Figure 2, Table 3). In the across-sex model (first column in Figure 2), predictive accuracy was moderate (*r* = 0.68), suggesting that anthropometric measurements serve as reasonable proxies for TIV. In contrast, the within-sex models (females: second and males: third columns in Figure 2) demonstrated a lower predictive accuracy (females: *r* = 0.26, males: *r* = 0.25), indicating the challenges of estimating TIV for each sex separately. This decrease in accuracy suggests that much of the predictive power in the across-sex model is driven by sexual dimorphism. Sex emerged as the most significant predictor in the across-sex model (Feature Importance; FI_sex_ = 71.05), reflecting differences in cranial capacity, which are likely influenced by hormonal factors, particularly testosterone, during development. Males, on average, exhibit 12% larger skulls and TIV compared to females, explaining the pronounced influence of sex on TIV prediction (Ruigrok et al. 2014; Eliot et al. 2021). However, body size reduces sex-based differences in TIV; when adjusted for height and weight, these differences are smaller, indicating TIV is linked to both body proportions and skull size. Age showed a smaller importance for TIV prediction (FI_age_: across-sex: 2.86, females: 0.56, males: 5.68), aligning with the stability of cranial structure across adulthood, unlike brain tissue volumes that decline with age-related atrophy. The slightly higher FI of age in males may reflect sex-specific changes in body composition or posture indirectly affecting TIV estimation. Among anthropometric predictors, seated height exhibited moderate positive associations with TIV (FI_seated height_: across-sex: 42.18, females: 29.72, males: 35.9), supporting a close link between vertical skeletal growth and cranial development. Weight also emerged as another important predictor (FI_weight_: across-sex: 26.2, females: 20.62, males: 27.65), likely due to its association with overall body size, though its relationship with TIV is weaker and more complex than that of height, potentially confounded by factors like adiposity or muscle mass. In addition, WC (FI_WC_: across-sex: −14.47, females: −8.77, males: −18.48) and HC (FI_HC_: across-sex: −9.88, females: −13.38, males: −6.64) displayed weak, negative associations with TIV across all models. These weak associations suggest that central adiposity has a limited influence on TIV. The results for the across-sex context using FreeSurfer, within-sex context, and RF model analysis are provided in the supplementary file (Figures S1, S2, S3).

**Table 3.** Feature importance (FI) of features.

| FI | TIV |  |  | TBV |  |  |
| --- | --- | --- | --- | --- | --- | --- |
|  | Both sexes | Female | Male | Both sexes | Female | Male |
| $FI_{\text{sex}}$ | 71.05 | -- | -- | 45.6 | -- | -- |
| $FI_{\text{age}}$ | 2.86 | 0.56 | 5.68 | -26.38 | -21.99 | -30.45 |
| $FI_{\text{weight}}$ | 26.2 | 20.62 | 27.65 | 21.31 | 16.03 | 22.26 |
| $FI_{\text{WC}}$ | -14.47 | -8.77 | -18.48 | -13.93 | -8.34 | -17.46 |
| $FI_{\text{HC}}$ | -9.88 | -13.38 | -6.64 | -5.63 | -7.7 | -3.55 |
| $FI_{\text{seated height}}$ | 42.18 | 29.72 | 35.9 | 35.15 | 26.06 | 29.49 |
| $FI_{\text{box height}}$ | -23.56 | -20.19 | -23.57 | -19.14 | -17.58 | -19.02 |

**Figure 2.**
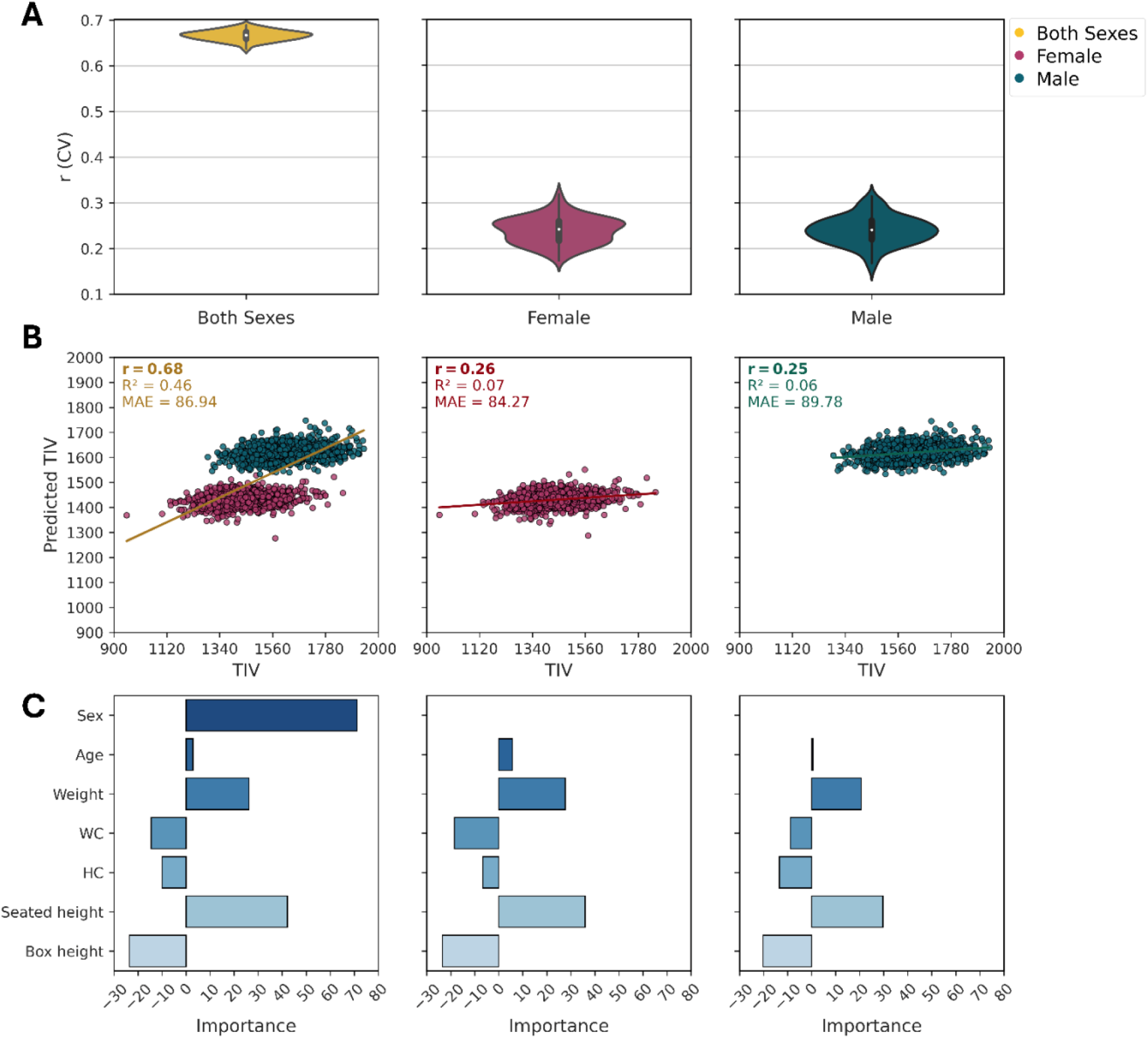
Estimation of head size (TIV) on CAT data using linear SVM. A) Prediction accuracy using the Pearson correlation coefficient (*r*) from CV analysis on the training dataset (90% of HC) for both sexes (left, *N* = 19,625), females (middle, *N* = 10,418), and male (right, *N* = 9,207). B) Association between predicted and true TIV values on the test dataset for both sexes (*N* = 2,182), females (*N* = 1,158), and males (*N* = 1,024). The scatter plot of predicted versus true TIV: for both sexes (*r* = 0.68, *R*^2^= 0.46, *MAE* = 86.94), for females (*r* = 0.26, *R*^2^= 0.07, *MAE* = 84.27), and for males (*r* = 0.25, *R*^2^= 0.06, *MAE* = 89.78). C) Feature importance (FI) scores from the final models trained on the training dataset (90% of HC). For both sexes (FI_Sex_ =71.05, FI_Age_ = 2.86, FI_Weight_ = 26.2, FI_WC_ = −14.47, FI_HC_ = −9.88, FI_Seated height_ = 42.18, FI_Box height_ = −23.56), for females (FI_Age_ = 0.56, FI_Weight_ = 20.62, FI_WC_ = −8.77, FI_HC_ = −13.38, FI_Seated height_ = 29.72, FI_Box height_ = −20.19), and for males (FI_Age_ = 5.68, FI_Weight_ = 27.65, FI_WC_ = −18.48, FI_HC_ = −6.64, FI_Seated height_ = 35.90, FI_Box height_ = −23.57).

#### 4.2.2. Prediction of brain size (TBV)

TBV estimation using the linear SVM model on CAT-derived data with age included as a feature was employed across both across-sex and within-sex cohorts (Figure 3, Table 4). In the across-sex model (first column in Figure 3), TBV prediction achieved moderate accuracy (*r* = 0.65), slightly lower than TIV (*r* = 0.68) but the within-sex models (Figure 3; females: second and males: third column) outperformed their TIV counterparts (females: *r* = 0.44, males: *r* = 0.45). This suggests that anthropometric measurements better capture brain size variation within in sex, unlike TIV, where sexual dimorphism dominates. Sex remained a significant predictor of TBV (FI_sex_ = 45.6), yet its influence was notably weaker than for TIV (FI_sex_ = 71.05). This reduced sexual dimorphism in TBV likely stems from the brain-to-cranium ratio: while males have larger skulls, their brain size does not scale proportionally with TIV (Ruigrok et al. 2014). Additionally, regional brain composition (e.g., such as females’ relatively higher GMV) may further moderate sex differences in TBV, making its relationship with body size more complex than for TIV (Ruigrok et al. 2014). A striking contrast with TIV emerged in the role of age. While TIV showed minimal age association, age was a strong negative predictor of TBV (FI_age_: across-sex: −26.38; females: −21.99; males: −30.45), indicating that older age is associated with lower TBV. This susceptibility to aging, potentially more pronounced in males, aligns with established patterns of tissue loss and further diminishes sexual dimorphism in TBV over time. Seated height and weight, like for TIV, remained important predictors of TBV. Seated height showed a strong positive predictor of TBV (FI_seated height:_ across-sex: 35.15, females: 26.06, males: 29.49), suggesting that, like TIV, taller individuals tend to have larger brain volumes. This positive relationship likely reflects a stronger connection to brain tissue development rather than cranial capacity, as TBV aligns more closely with neurodevelopmental scaling, whereas TIV is influenced by broader skeletal factors. Thus, seated height may be a more direct predictor of brain tissue volume. However, the predictive power of weight (FI_weight_: across-sex: 21.31, females: 16.03, males: 22.26) was somewhat less pronounced than for TIV, indicating that body size may have a more complex relationship with brain size due to factors like body composition, such as fat and muscle distribution, which differs between sexes. WC and HC also showed negative association with TBV (FI_WC_ = −13.93, females: −8.34, males: −17.46; FI_HC_ = −5.63, females: - 7.7, males: −3.55). However, TBV negative associations were slightly weaker in magnitude and more variable across sexes, possibly due to tissue-specific factors such as age-related GMV loss, metabolic effects of adiposity, and sex differences in brain composition. These factors may influence TBV more heterogeneously than the fixed cranial structure of TIV. The results for the across-sex context using FreeSurfer, within-sex context, and RF model analysis are provided in the supplementary file (Figures S4, S5, S6).

**Figure 3.**
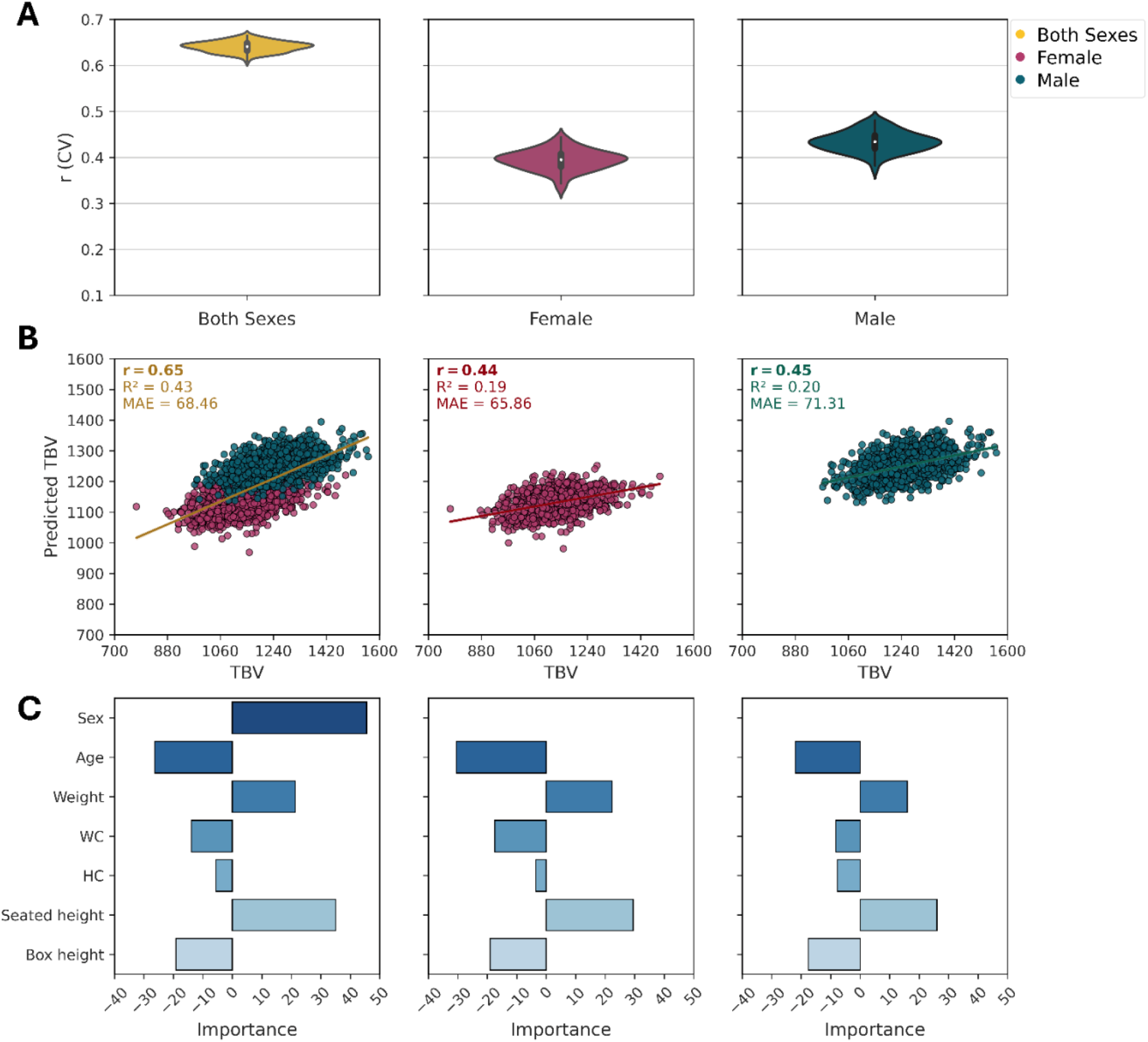
Estimation of brain size (TBV) on CAT data using linear SVM. A) Prediction accuracy using the Pearson correlation coefficient (*r*) from CV analysis on the training dataset (90% of HC) for both sexes (left, *N* = 19,625), females (middle, *N* = 10,418), and male (right, *N* = 9,207). B) Association between predicted and true TIV values on the test dataset for both sexes (*N* = 2,182), females (*N* = 1,158), and males (*N* = 1,024). The scatter plot of predicted versus true TIV: for both sexes (*r* = 0.68, *R*^2^= 0.46, *MAE* = 86.94), for females (*r* = 0.26, *R*^2^= 0.07, *MAE* = 84.27), and for males (*r* = 0.25, *R*^2^= 0.06, *MAE* = 89.78). C) Feature importance (FI) scores from the final models trained on the training dataset (90% of HC). For both sexes (FI_Sex_ = 45.6, FI_Age_ = −26.38, FI_Weight_ = 21.31, FI_WC_ = −13.93, FI_HC_ = −5.63, FI_Seated height_ = 35.15, FI_Box height_ = - 19.14), for females (FI_Age_ = −21.99, FI_Weight_ = 16.03, FI_WC_ = −8.34, FI_HC_ = −7.7, FI_Seated height_ = 26.06, FI_Box height_ = −17.58), and for males (FI_Age_ = −30.45, FI_Weight_ = 22.26, FI_WC_ = −17.46, FI_HC_ = −3.55, FI_Seated height_ = 29.49, FI_Box height_ = −19.02).

**Figure 4.**
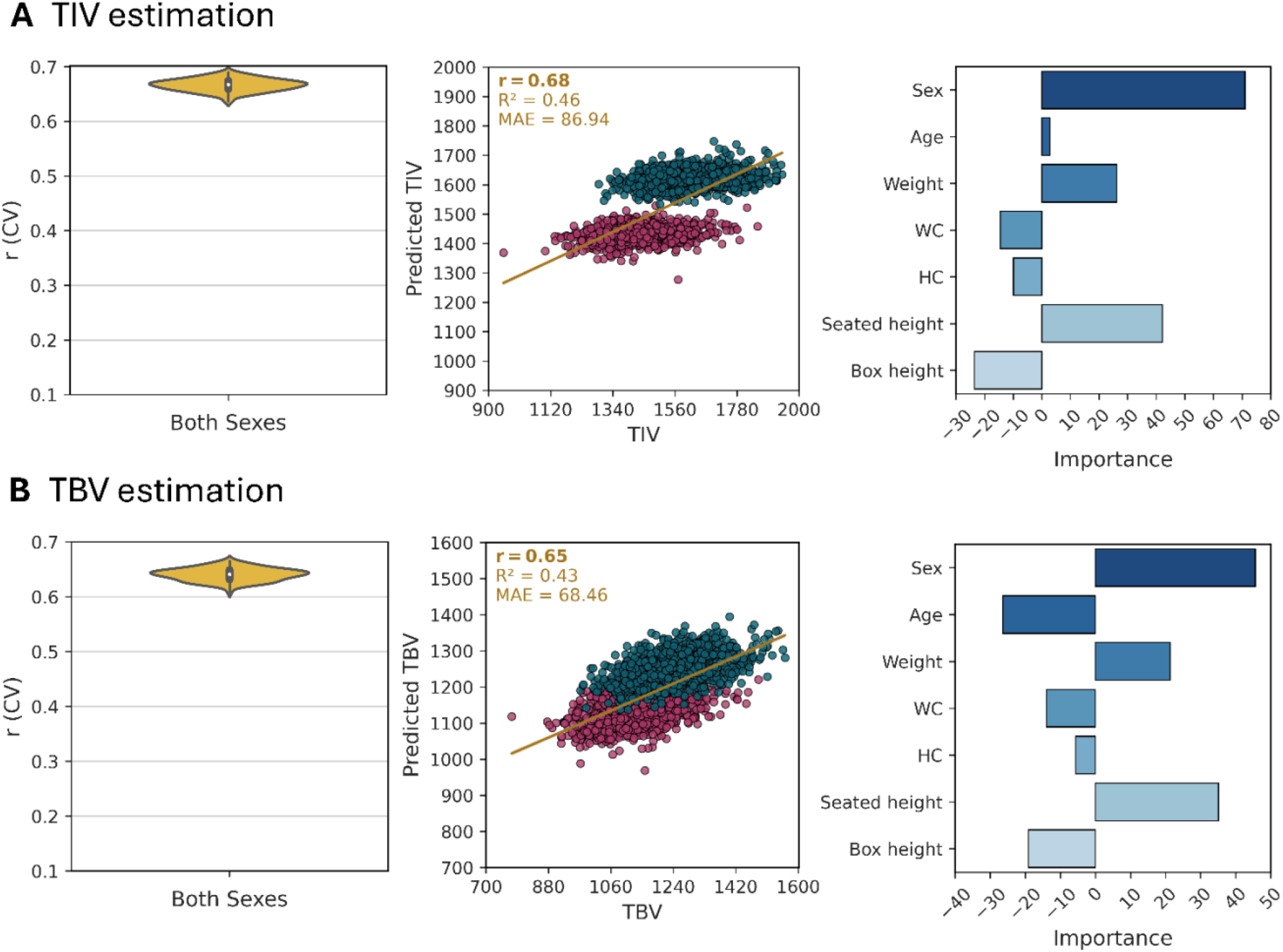
Comparison between head size and brain size including standing height as a feature. A) Estimation of head size (TIV) and B) Estimation of brain size (TBV) on CAT data using linear SVM.

**Figure 5.**
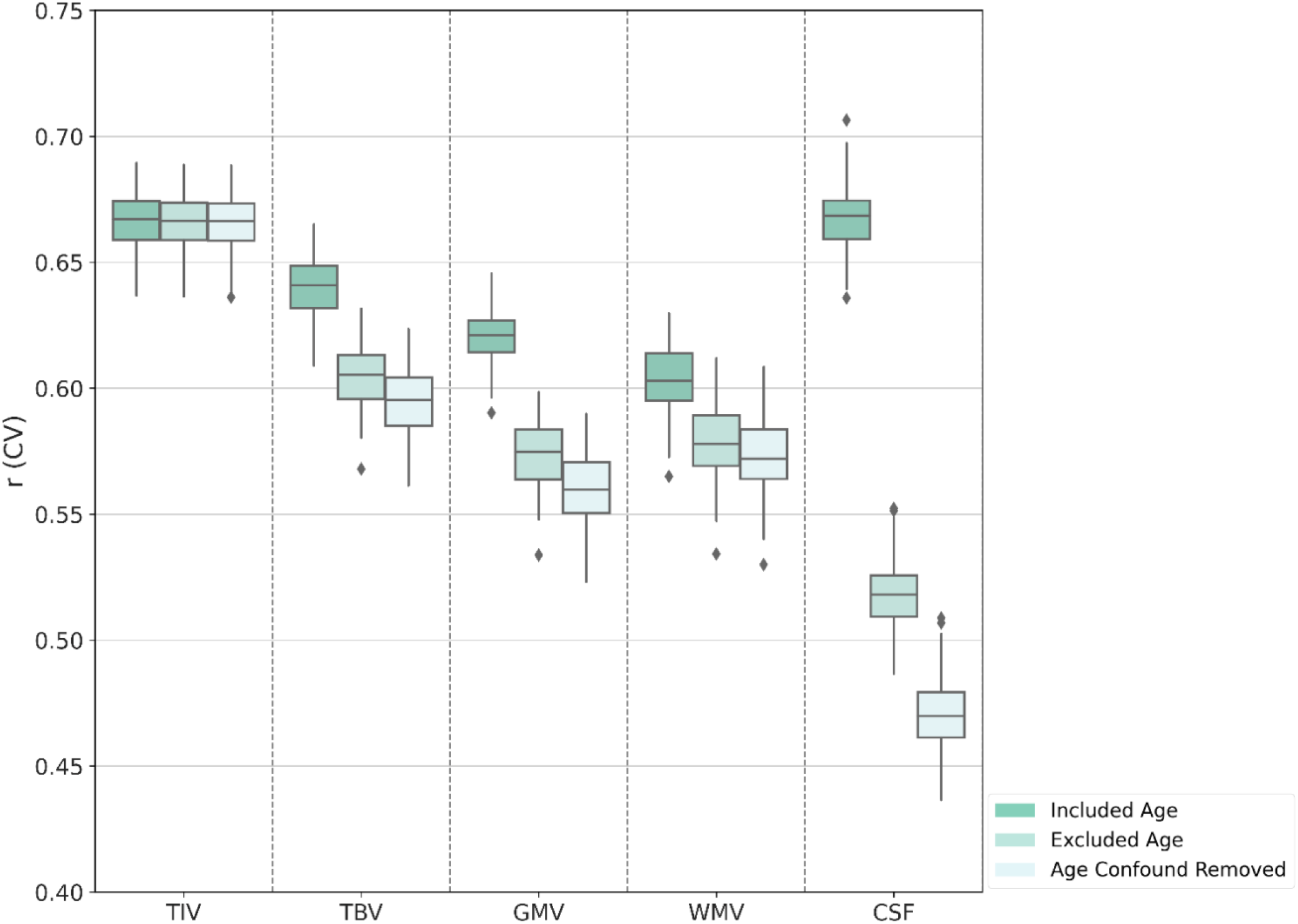
Impact of age on brain volumes on CAT data using linear SVM. Comparison of brain volumes concerning age-related effects across different age-related features. The analysis includes three conditions: including age as a feature (Included Age), where age is included as a predictor in the model; excluding age from the feature set (Excluded Age), where age is omitted from the model as a predictor, and impact of age after confounder removal (Age Confound Removed), where age-related effects are assessed following the removal of potential confounding factors associated with age.

### 4.3. Comparison between standing height and seated height

The comparison between seated height and standing height is crucial for understanding the complex relationships between body measurements and brain morphology (Masanovic et al. 2020). Standing height includes both trunk and leg length, with the femur significantly contributing to overall height. In contrast, seated height isolates trunk growth, providing a more direct assessment of anatomical factors that might be related to brain size. This distinction is important due to the differing growth patterns of the femur compared to the trunk and skull, which can introduce confounding variables when using standing height as a predictor of brain size. Therefore, in this analysis, we compared the predictive power of standing height and seated height for the head size (TIV) and brain size (TBV) by including standing height as an additional feature (Figure 4). Standing height exhibited a higher predictive power than seated height for both TIV (FI_standing height_ = 23.37, FI_seated height_ = 10.58) and TBV (FI_standing height_ = 16.51, FI_seated height_ = 11.56) prediction. These high FI scores suggest that standing height captures greater variability in the brain volumes, likely due to its inclusion of leg length. However, despite these higher FI scores, the addition of standing height did not improve overall prediction accuracy (TIV: *r* = 0.68, TBV: *r* = 0.65). This indicates that while standing height captures more variability, it does not provide additional predictive power beyond what seated height already offers. The lack of improvement in prediction accuracy, despite higher FI scores for standing height, can be explained by developmental and anatomical factors. The trunk and skull, which are reflected in seated height, follow a growth trajectory that aligns more closely with brain development, reaching near-adult proportions earlier than appendicular structures like the femur. In contrast, leg length, a significant component of standing height, exhibits more prolonged and variable growth may be influenced by environmental, nutritional, or postural factors. As a result, seated height, focusing on the upper body, may serve as a anatomically relevant proxy for cranial and brain size, potentially minimizing the noise introduced by leg length variability. Thus, while standing height captures greater variability as reflected in its higher FI scores, it does not translate into better predictive performance, reinforcing the idea that trunk-based measures more closely capture biologically relevant scaling between body and brain size, reflected by TIV and TBV.

### 4.4. Impact of age on brain volumes

The comparison of brain volume predictions across three age strategies—including age as a feature, excluding it from the feature set, or removing it as a confound (see Section 3.1)—provides valuable insights into how age influences various brain volumes, particularly when considering anthropometric features. The results were obtained using CAT-derived data in an across-sex context (Figure 5). The analysis highlights the intricate relationship between age, brain structure, and anthropometric measurements. Distinct age-related patterns emerged in brain volume predictions, emphasizing the complex interplay of these factors. TIV and TBV predictions showed distinct patterns with age. TIV predictions remained remarkably stable (*r* ≈ 0.66) across all configurations, indicating a minimal impact of age. This stability aligns with the understanding that, on a population level, TIV remains fixed throughout adulthood, largely due to the stable nature of the bony structure of the cranium, which maintains its size despite age-related brain atrophy (Smith et al. 2007). The consistent prediction accuracy suggests that TIV is primarily influenced by genetic or developmental factors, rather than age or anthropometrics. In contrast, TBV exhibited marked sensitivity to age. When age was included as a feature in the prediction model, the accuracy of TBV prediction (*r* = 0.64) reflected the strong relationship between age and changes in GMV and WMV over the lifespan. As individuals age, GMV generally decreases, and WMV may also experience a decline, both of which contribute to overall reductions in TBV (Mills et al. 2016; Irimia 2021; Yamada et al. 2023). In this context, the inclusion of age as a feature captured not only these structural changes but also provided critical insight into how TBV changes over time. Excluding age from the feature set or removing it as a confound led to decreased accuracy (*r* = 0.6 and *r* = 0.59, respectively), confirming that age is a critical factor in TBV prediction, even when accounting for other variables like anthropometric measurements or sex. The similar accuracies in the age-excluded and age-confound-removed conditions suggest that age-related effects on TBV are primarily direct rather than merely confounding influences. This direct effect of age on TBV highlights how aging affects the brain overall volume in a measurable, predictable way, unlike TIV, which remains stable.

GMV and WMV both showed reduced prediction accuracies when age was excluded from the feature set or removed as a confounder, with a more pronounced impact for GMV. For GMV, the prediction accuracy dropped when age was excluded from the feature set (*r* = 0.57) or removed as a confound (*r* = 0.55), compared to when age was included as a feature (*r* = 0.62). These reductions indicate that age plays a critical role in predicting GMV beyond what can be accounted for by anthropometric features. This finding is consistent with the understanding that GMV atrophy is more pronounced with age due to processes such as neuronal loss, dendritic pruning, and synaptic changes, which occur more significantly in certain brain regions (Gennatas et al. 2017; Irimia 2021; Yamada et al. 2023). Age, therefore, serves as a proxy for these ongoing structural changes, and removing it as a confounder may mask important age-specific variations in GMV. This loss of predictive accuracy suggests that age-related changes in brain structure are not entirely captured by other variables, such as anthropometric features, implying the importance of age as a crucial marker for predicting GMV changes (Gennatas et al. 2017). In contrast, the impact of age on WMV prediction was less pronounced, though still significant. Prediction accuracy dropped moderately when age was excluded from feature set (*r* = 0.58) or removed as a confound (*r* = 0.57), compared to when age was included as a feature (*r* = 0.6). This suggests that, while anthropometric features are important to WMV prediction, age still adds predictive value. The modest decrease in prediction accuracy when age is excluded or removed as a confound suggests that anthropometric features may be more strongly associated with WMV than with GMV. The minimal difference between the age-excluded and age-confound removed conditions suggest that age-related effects on WMV are direct, with limited confounding influences from other predictors. Interestingly, these findings suggest that age-related atrophy is more pronounced in GMV than in WMV, which is consistent with the established pattern of brain shrinkage in normal aging (Smith et al. 2007; Irimia 2021). GMV atrophy tends to occur earlier and more extensively than WMV loss, which likely explain the stronger impact of age on GMV predictions. However, this differential atrophy could also be influenced by segmentation bias. Variations in the methods used to segment GMV and WMV could contribute to the observed differences in volume and prediction accuracy (Smith et al. 2007; Irimia 2021).

For the CSF volume, age had a strong impact on prediction accuracy. When age was excluded from the feature set or removed as a covariate, accuracy dropped substantially (*r* = 0.51, *r* = 0.46, respectively) compared to when age was included in the feature set (*r* = 0.66). These large differences highlight that age is a critical factor in predicting CSF volume and are consistent with previous research showing an approximate 30ml (2%) increase per decade from the 20s to 488ml (33.7%) in individuals over 80 years old, due to tissue loss and fluid expansion with aging (Yamada et al. 2023). The further reduction in accuracy when age was removed as a confounder suggests that age-related effects on CSF volume prediction involve both direct and indirect contributions. The direct effect refers to the structural changes, particularly the atrophy of GMV and WMV, which leads to an increase in CSF volume. The indirect effect include other factors, such as anthropometric features and sex, that can influence brain structure and the relationship between age and CSF. This is consistent with the known relationship between GMV and WMV atrophy and the increase in CSF volume with age (Gennatas et al. 2017; Irimia 2021; Yamada et al. 2023). As both GMV and WMV decrease, CSF volume increases with aging, providing a clearer and more consistent indicator of age-related structural changes. These findings underscore the importance of including age as a factor in neuroimaging studies, particularly when predicting brain volumes, as it captures both direct and indirect age-related changes in brain structure. The results for the across-sex context using FreeSurfer, within-sex context, and RF model analysis are provided in the supplementary file (Figures S7, S8, S9, S10, S11, S12, and S13). These additional analyses corroborate the main findings, supporting the consistency and robustness of the results across different methods and contexts.

## 5. Conclusions

This study provides a comprehensive analysis of how brain volumes are associated with anthropometric features, age, and sex, in a large UK Biobank cohort using advanced machine learning methods. Our findings reveal that TIV is predominantly driven by sex, reflecting a strong sexual dimorphism in cranial development. In contrast, TBV, GMV, WMV, and CSF are more sensitive to age, consistent with well-established patterns of brain tissue loss and compensatory fluid expansion across adulthood, with the greatest age-related sensitivity observed in CSF and GMV. Anthropometric measures, including weight, height, waist circumference, and hip circumference, provide additional predictive value, particularly for TBV and TIV. However, their influences are secondary to those of age and sex. Notably, central adiposity (as indexed by waist circumference) is negatively associated with brain tissue volumes, likely reflecting the metabolic impact of visceral fat, while height and weight are positively linked to cranial and brain size, especially earlier in life. However, these relationships are complex and can vary across brain volumes and demographic groups. The differential association of age, sex, and anthropometric features across brain compartments highlights the necessity of including both age and sex as core covariates in neuroimaging research to ensure accurate interpretation of volumetric differences. These results also underscore the value of integrating detailed anthropometric data to refine models of brain structure variation and to better understand the biological and environmental factors underlying brain morphology. From a clinical perspective, these findings may aid the interpretation of brain imaging by helping distinguish normative, age-related brain volume changes from neurodegenerative disease. Overall, this work advances our understanding of the factors driving variation in brain volumes and provides a foundation for future longitudinal studies to unravel the mechanisms linking physical development, aging, and neuroanatomy. Such insights are crucial for interpreting neuroimaging findings in both healthy and clinical populations and for informing strategies aimed at promoting brain health across the lifespan.

## Supplementary Information

The online version contains supplementary material available at supplementary.docx.

## Acknowledgments

This research has been conducted using data from UK Biobank resources (application number 41655). All data used in this study are publicly accessible from UK Biobank via their standard data access procedure (http://www.ukbiobank.ac.uk/).

## Author contributions

All authors read and approved the final manuscript. We use the CRediT contributor role taxonomy to describe individual contributions to the paper. Conceptualization: K.N., S.B.E., C.G., K.R.P.; Data curation: K.N., V.K., K.R.P.; Formal analysis: K.N.; Methodology: K.N., S.B.E., C.G., K.R.P.; Supervision: C.G., S.B.E., K.R.P.; Visualization: K.N., Writing—original draft: K.N.; Writing—review & editing: K.N., S.B.E., G.A., L.H., C.T., V.K., F.R., C.G., K.R.P.

## Ethics approval and consent to participate

The UK Biobank study was approved by the North West Multicenter Research Ethics Committee (No. 16/NW/0274), with written informed consent obtained from all participants. A re-analysis of the anonymized data was approved by the ethics committee of the Heinrich Heine University Düsseldorf (2018-317-RetroDEuA).

## Data availability

All data used in this study are publicly available through the UK Biobank, accessible via their standard data access procedure at http://www.ukbiobank.ac.uk/. The code used in the current study is available from the authors upon reasonable request.

