## Supplementary Material for "Predicting Brain Volumes from Anthropometric and Demographic Features: Insights from UK Biobank Neuroimaging Data"

**List of tables and figures**

| **Table S1** | Definition of the healthy controls (HC) based on ICD-10 exclusion criteria |
| --- | --- |
| **Table S2** | Comparison of linear SVM and RF models on the training dataset for all brain volumes |
| **Figure S1** | Prediction of head size (TIV) on CAT data using RF. |
| **Figure S2** | Prediction of head size (TIV) on FreeSurfer data using linear SVM. |
| **Figure S3** | Prediction of head size (TIV) on FreeSurfer data using RF. |
| **Figure S4** | Prediction of brain size (TBV) on CAT data using RF. |
| **Figure S5** | Prediction of brain size (TBV) on FreeSurfer data using linear SVM. |
| **Figure S6** | Prediction of brain size (TBV) on FreeSurfer data using RF. |
| **Figure S7** | Impact of age on brain volumes for across-sex analysis on CAT data using RF. |
| **Figure S8** | Impact of age on brain volumes for within-sex analysis on CAT data using linear SVM. |
| **Figure S9** | Impact of age on brain volumes for within -sex analysis on CAT data using RF. |
| **Figure S10** | Impact of age on brain volumes for across-sex analysis on FreeSurfer data using linear SVM. |
| **Figure S11** | Impact of age on brain volumes for across -sex analysis on FreeSurfer data using RF. |
| **Figure S12** | Impact of age on brain volumes for within-sex analysis on FreeSurfer data using linear SVM. |
| **Figure S13** | Impact of age on brain volumes for within-sex analysis on FreeSurfer data using RF. |

**Table S1:** Definition of the healthy controls (HC) based on ICD-10 exclusion criteria

| Populations | Excluded ICD-10 criteria |
| --- | --- |
| Healthy controls (HC) | Mental and behavioural disorders: F |
|  | Diseases of the nervous system: G |
|  | Cerebrovascular diseases: I60-I69 |
|  | Diseases of the musculoskeletal system and connective tissue: M |
|  | Injury, poisoning, and certain other consequences of external causes: S |

**Table S2:** Comparison of linear SVM and RF models on the training dataset for all brain volumes in CAT and FreeSurfer data.

|  |  |  |  | Linear SVM | | |  | RF | | |
| --- | --- | --- | --- | --- | --- | --- | --- | --- | --- | --- |
| Data | Brain volume | Scores (median) |  | Both sexes | Female | Male |  | Both sexes | Female | Male |
| CAT | TIV | Pearson’s r |  | 0.67 | 0.24 | 0.24 |  | 0.63 | 0.16 | 0.16 |
|  |  | R2 |  | 0.44 | 0.06 | 0.06 |  | 0.4 | -0.02 | -0.02 |
|  |  | MAE |  | 87.98 | 82.24 | 94.81 |  | 91.64 | 85.64 | 98.0 |
|  | TBV | Pearson’s r |  | 0.64 | 0.4 | 0.43 |  | 0.61 | 0.33 | 0.38 |
|  |  | R2 |  | 0.41 | 0.16 | 0.19 |  | 0.36 | 0.08 | 0.12 |
|  |  | MAE |  | 70.71 | 66.03 | 75.52 |  | 73.49 | 68.74 | 78.58 |
|  | GMV | Pearson’s r |  | 0.62 | 0.41 | 0.45 |  | 0.59 | 0.35 | 0.39 |
|  |  | R2 |  | 0.39 | 0.17 | 0.2 |  | 0.34 | 0.1 | 0.13 |
|  |  | MAE |  | 36.41 | 34.33 | 38.6 |  | 37.93 | 35.87 | 40.3 |
|  | WMV | Pearson’s r |  | 0.6 | 0.33 | 0.37 |  | 0.57 | 0.25 | 0.31 |
|  |  | R2 |  | 0.36 | 0.11 | 0.13 |  | 0.31 | 0.03 | 0.07 |
|  |  | MAE |  | 40.15 | 37.31 | 43.24 |  | 41.64 | 38.92 | 44.69 |
|  | CSF | Pearson’s r |  | 0.67 | 0.43 | 0.57 |  | 0.64 | 0.36 | 0.53 |
|  |  | R2 |  | 0.45 | 0.18 | 0.33 |  | 0.41 | 0.1 | 0.27 |
|  |  | MAE |  | 38.24 | 36.78 | 39.22 |  | 39.35 | 38.27 | 40.62 |
| FreeSurfer | TIV | Pearson’s r |  | 0.62 | 0.3 | 0.25 |  | 0.58 | 0.22 | 0.18 |
|  |  | R2 |  | 0.38 | 0.09 | 0.06 |  | 0.33 | 0.01 | -0.01 |
|  |  | MAE |  | 95048.98 | 87560.94 | 103153.57 |  | 98794.95 | 91282.96 | 106920.35 |
|  | TBV | Pearson’s r |  | 0.64 | 0.42 | 0.4 |  | 0.6 | 0.35 | 0.35 |
|  |  | R2 |  | 0.41 | 0.17 | 0.16 |  | 0.35 | 0.1 | 0.1 |
|  |  | MAE |  | 68753.22 | 64813.69 | 73012.3 |  | 71575.78 | 67599.69 | 75485.72 |
|  | GMV | Pearson’s r |  | 0.62 | 0.5 | 0.48 |  | 0.58 | 0.45 | 0.43 |
|  |  | R2 |  | 0.38 | 0.24 | 0.23 |  | 0.33 | 0.18 | 0.17 |
|  |  | MAE |  | 34660.01 | 32960.94 | 36487.76 |  | 36145.6 | 34363.87 | 38102.41 |
|  | WMV | Pearson’s r |  | 0.62 | 0.3 | 0.3 |  | 0.58 | 0.22 | 0.24 |
|  |  | R2 |  | 0.38 | 0.09 | 0.09 |  | 0.34 | 0.01 | 0.03 |
|  |  | MAE |  | 38691.92 | 36234.39 | 41546.71 |  | 40307.91 | 37815.13 | 42878.44 |
|  | CSF | Pearson’s r |  | 0.43 | 0.23 | 0.24 |  | 0.37 | 0.15 | 0.16 |
|  |  | R2 |  | 0.18 | 0.05 | 0.06 |  | 0.11 | -0.03 | -0.03 |
|  |  | MAE |  | 182.61 | 168.89 | 198.03 |  | 191.3 | 176.41 | 207.13 |

**Figure S1:** Prediction of head size (TIV) on CAT data using RF.

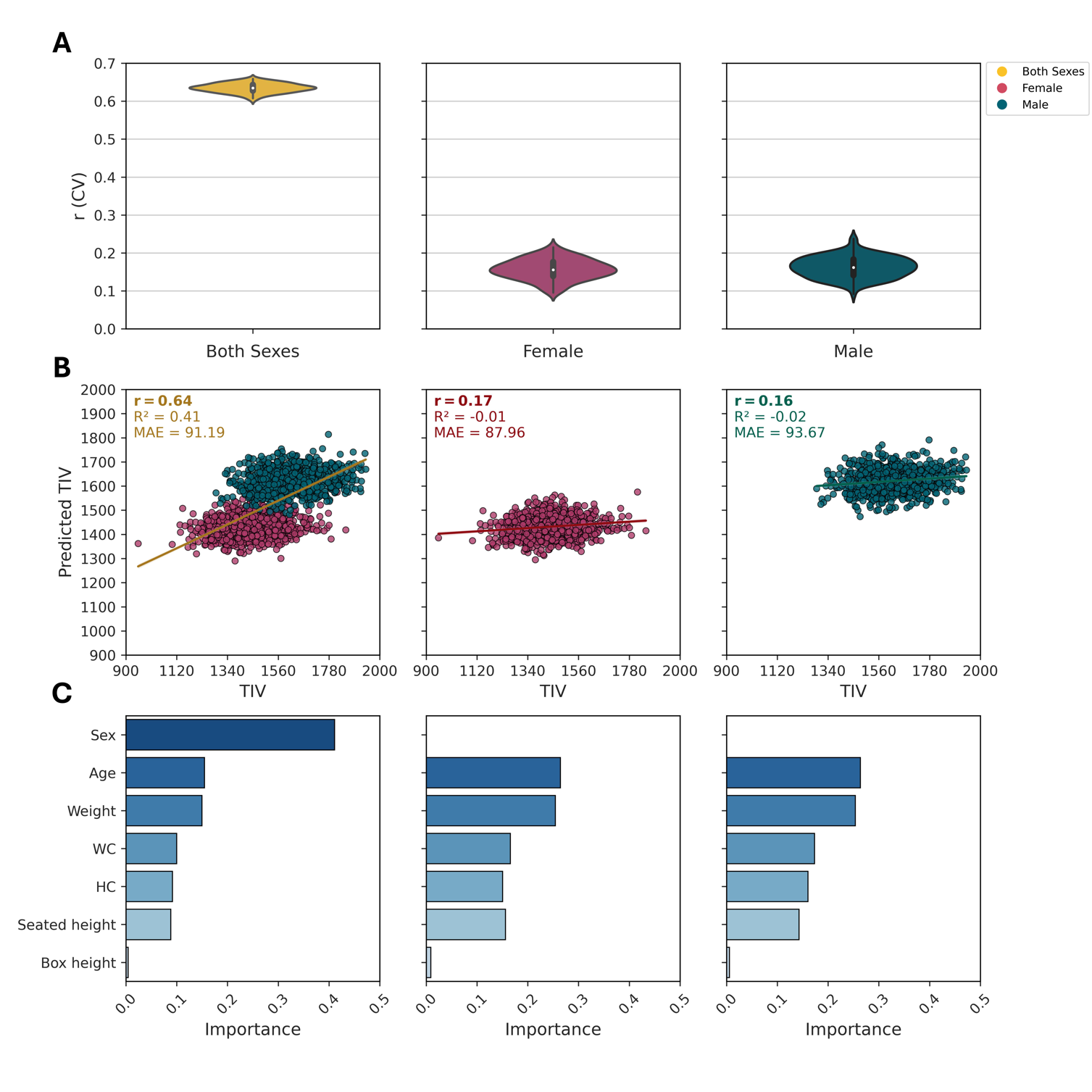

**Figure S2:** Prediction of head size (TIV) on FreeSurfer data using linear SVM.

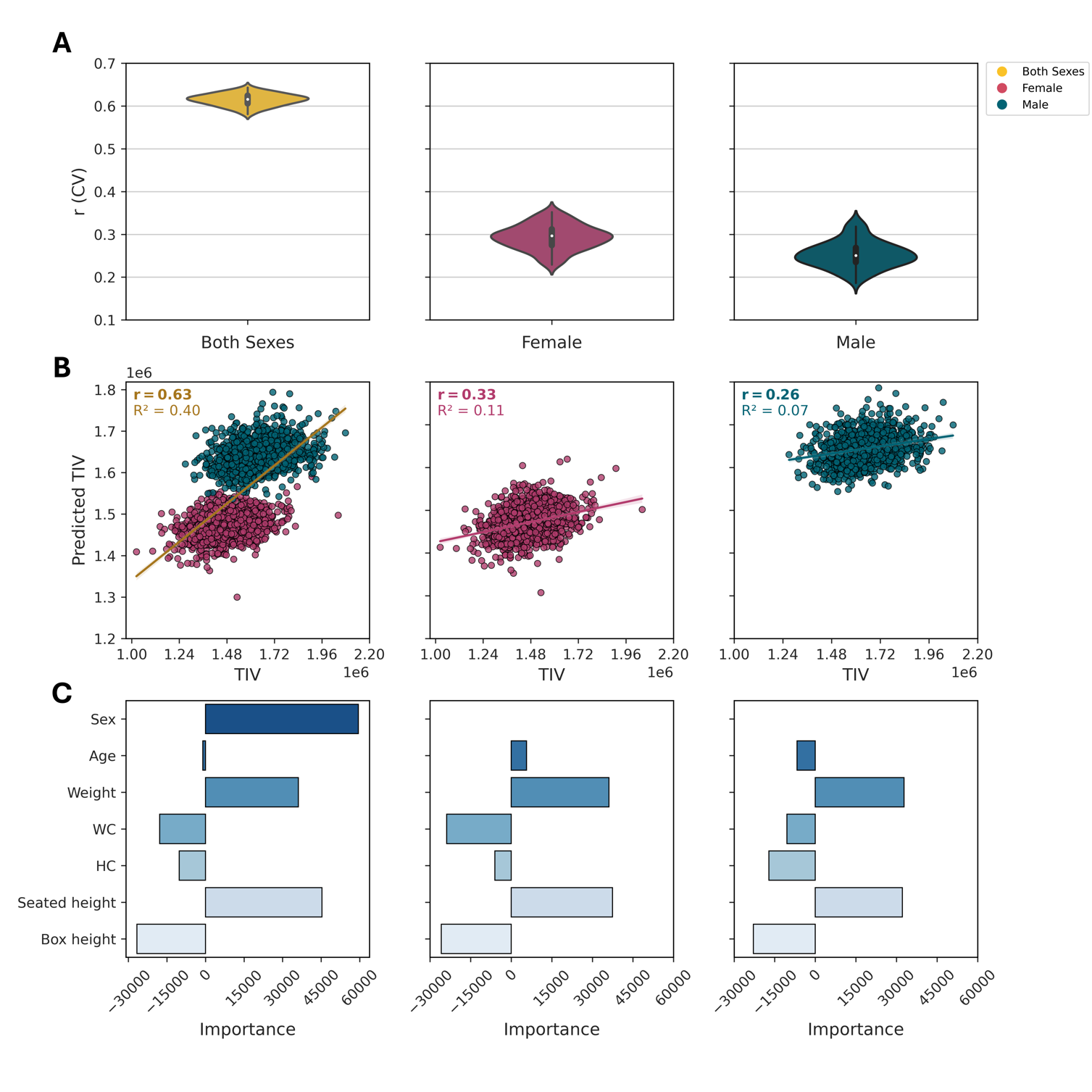

**Figure S3:** Prediction of head size (TIV) on FreeSurfer data using RF.

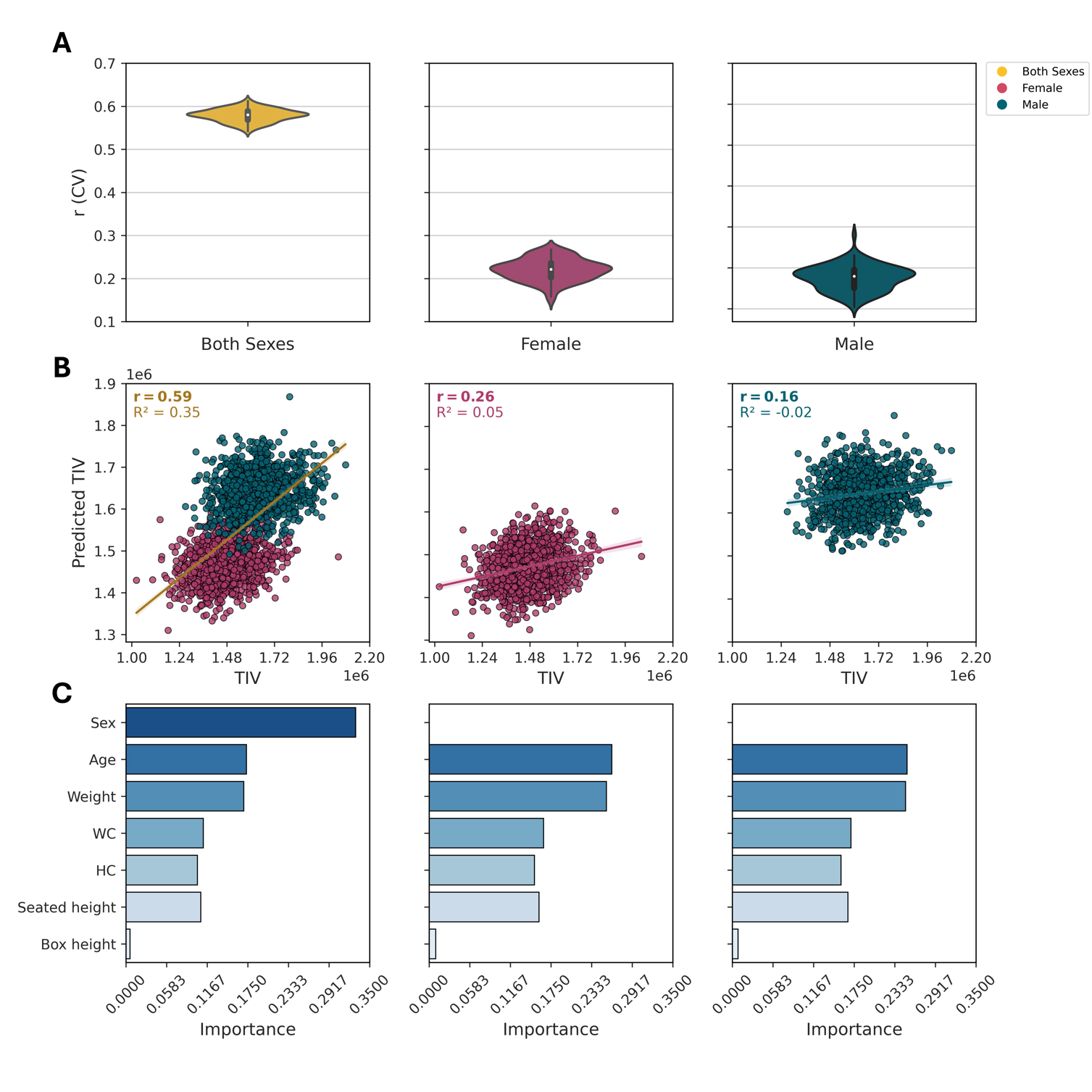

**Figure S4:** Prediction of brain size (TBV) on CAT data using RF.

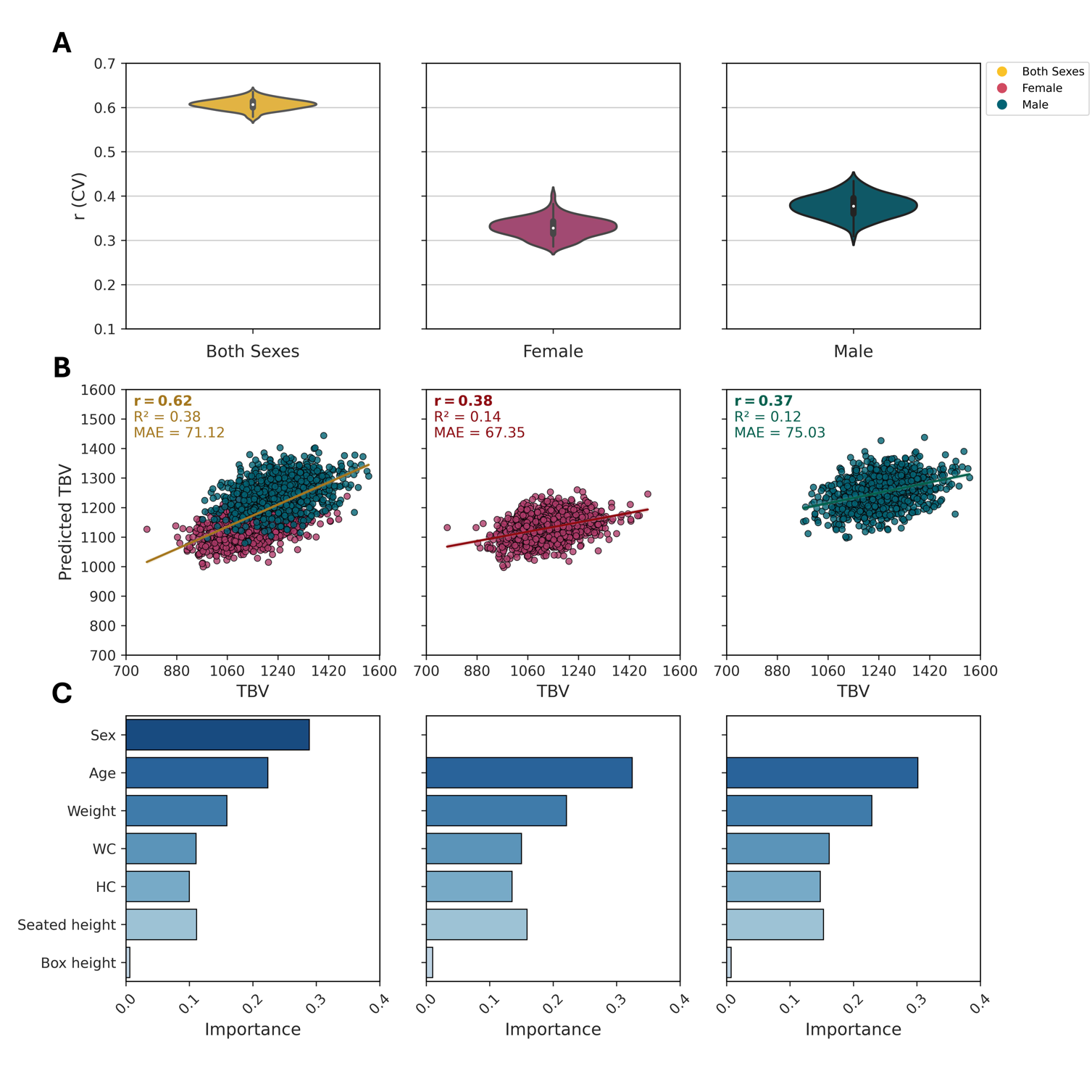

**Figure S5:** Prediction of brain size (TBV) on FreeSurfer data using linear SVM

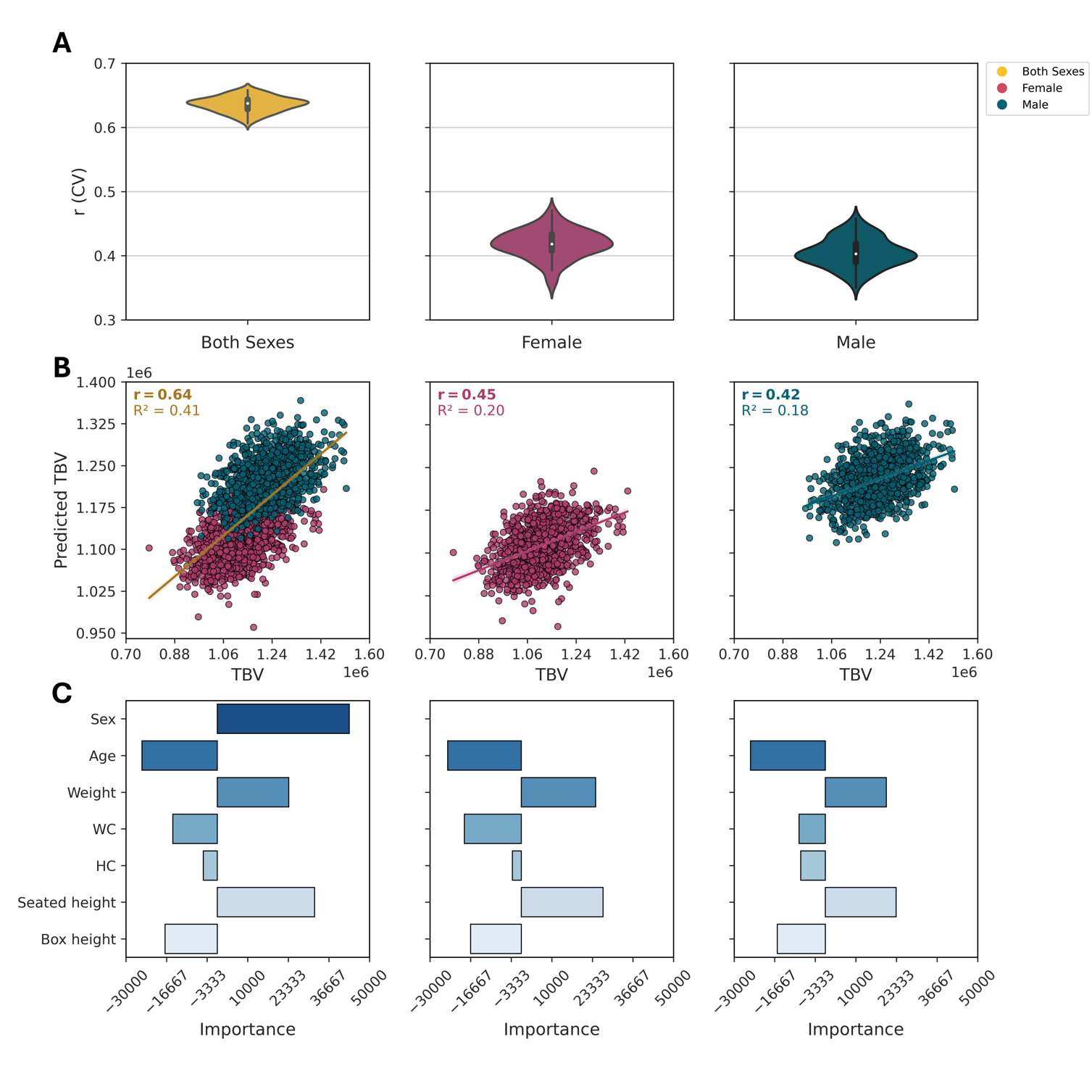

**Figure S6:** Prediction of brain size (TBV) on FreeSurfer data using RF.

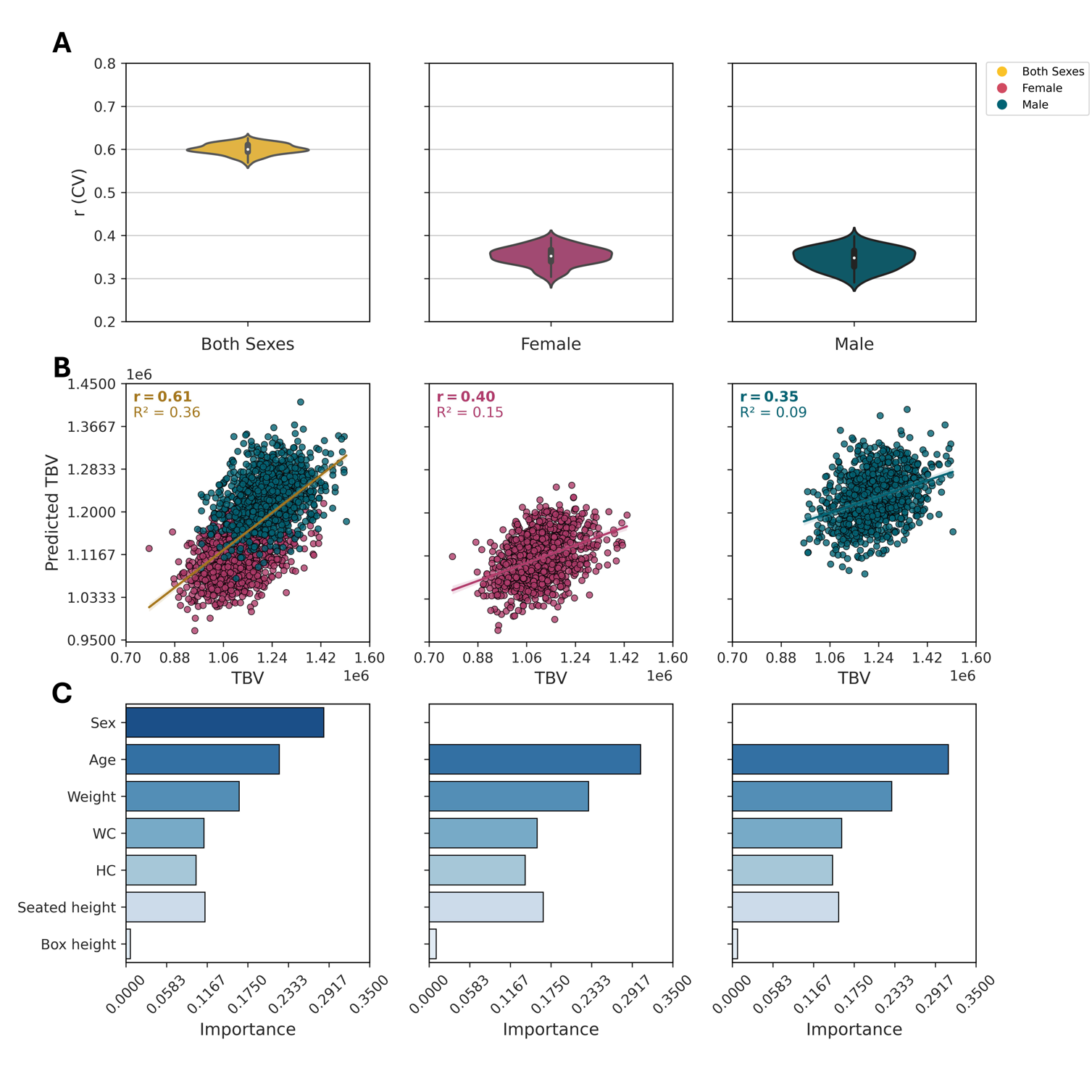

**Figure S7:** Impact of age on brain volumes for across-sex analysis on CAT data using RF.

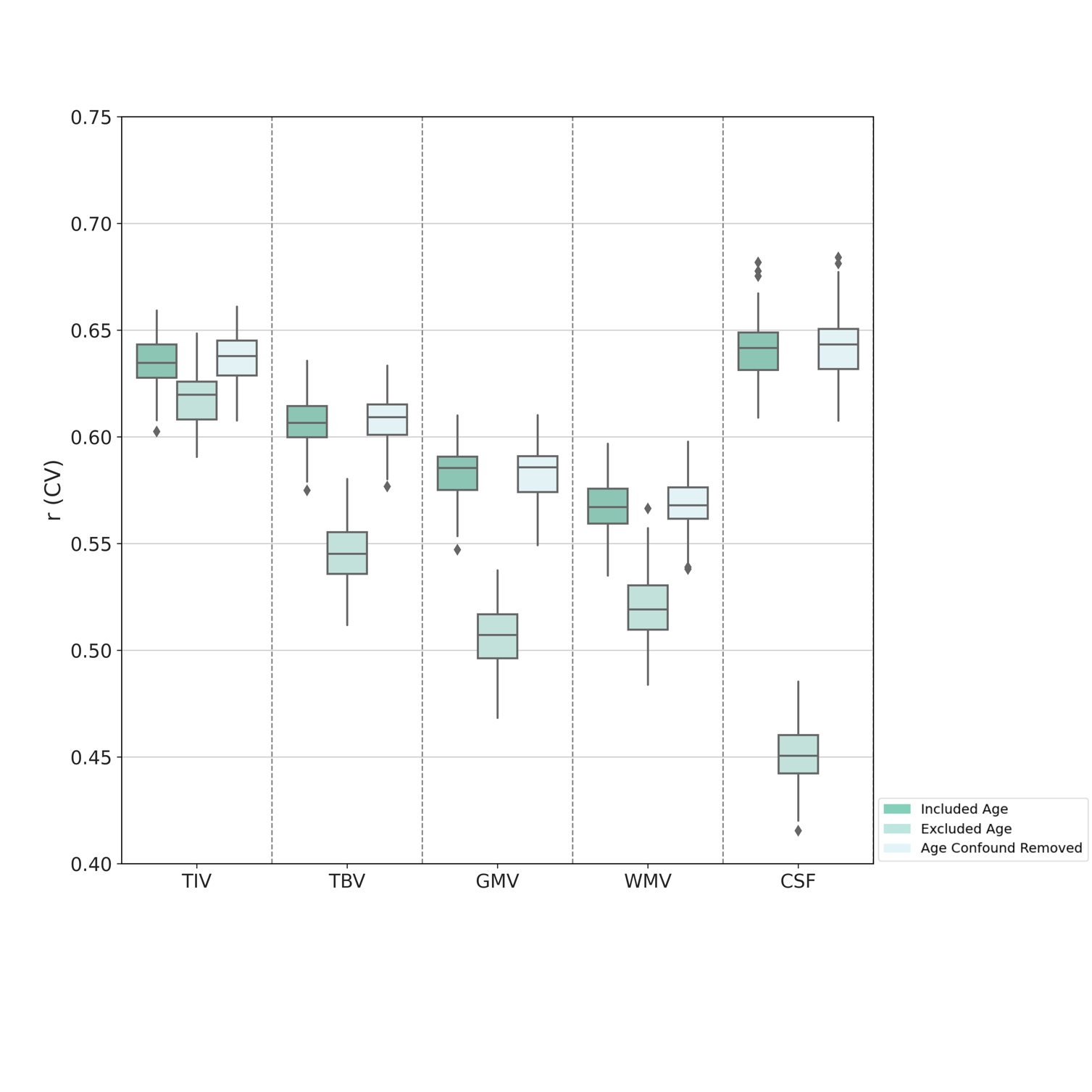

**Figure S8:** Impact of age on brain volumes for within-sex analysis on CAT data using linear SVM.

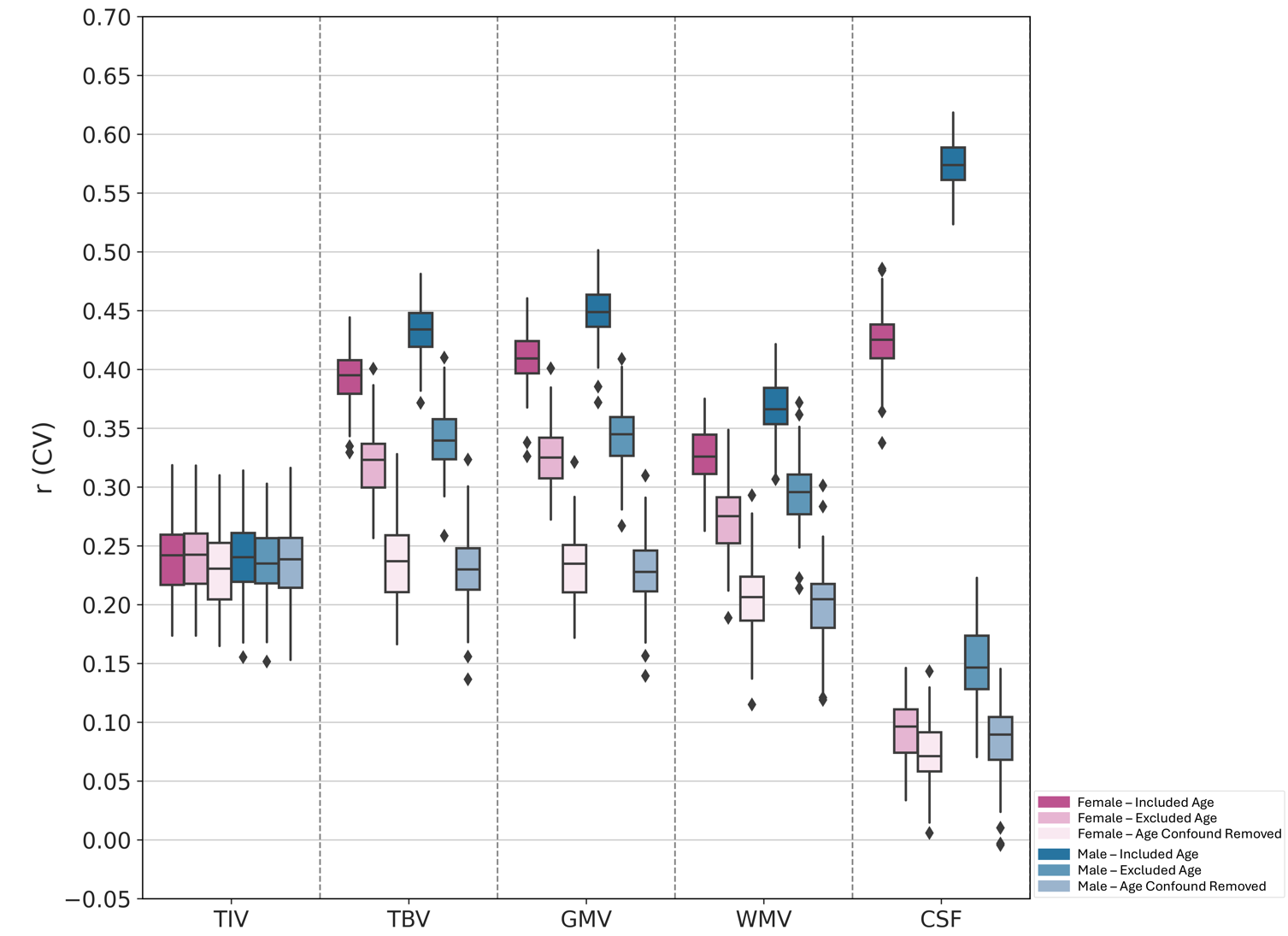

**Figure S9:** Impact of age on brain volumes for within-sex analysis on CAT data using random forest.

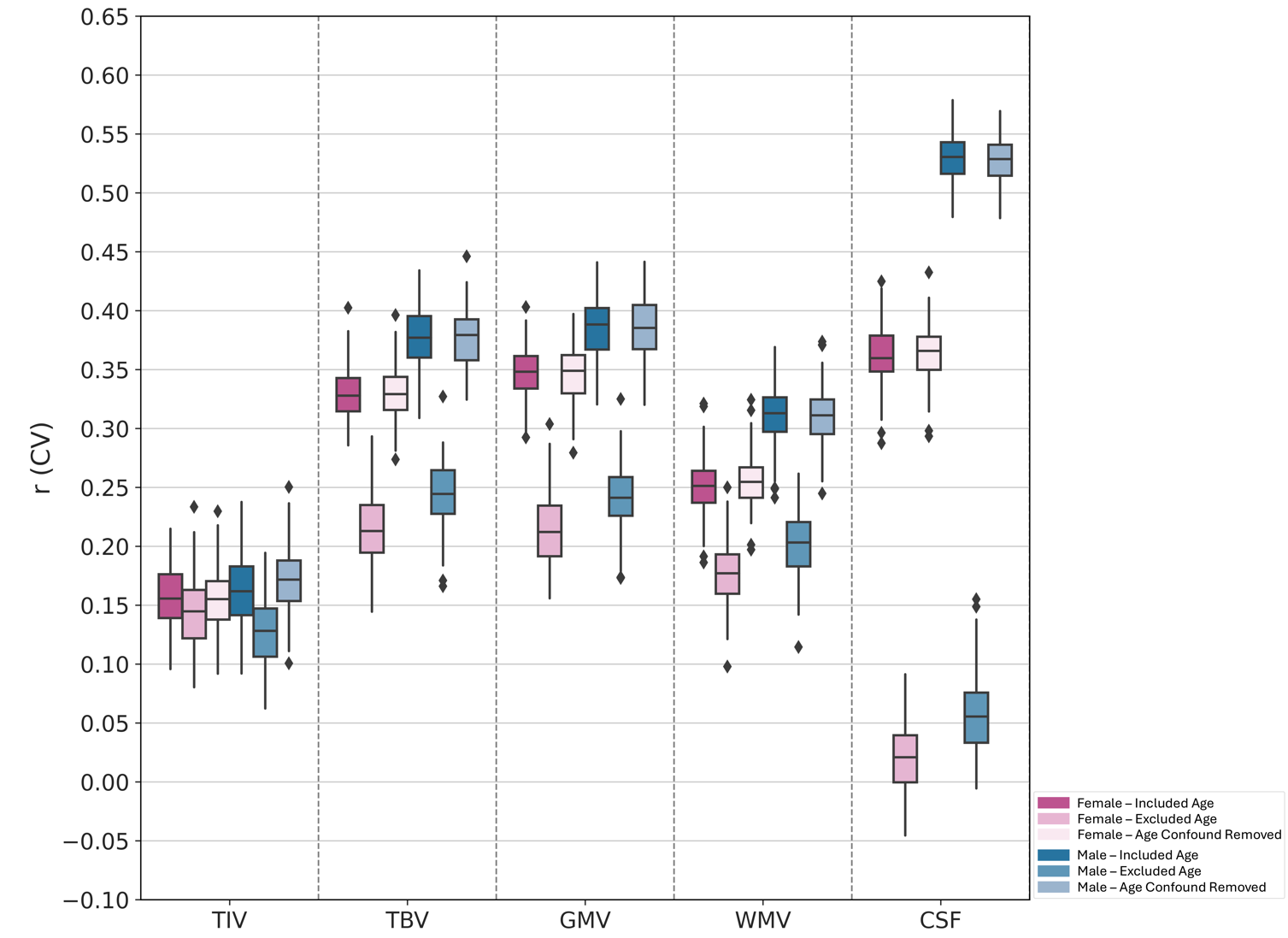

**Figure S10:** Impact of age on brain volumes for across-sex analysis on FreeSurfer data using linear SVM.

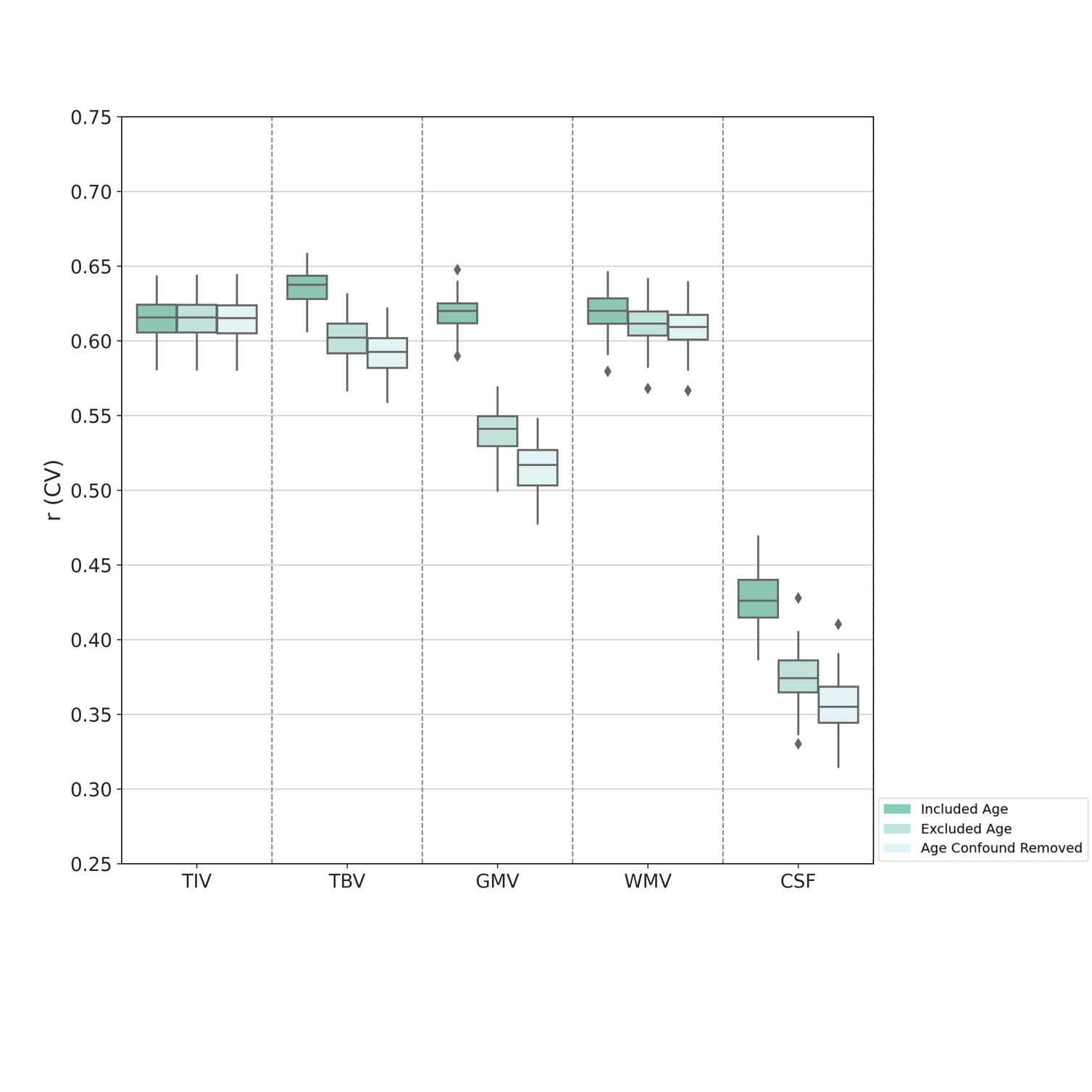

**Figure S11:** Impact of age on brain volumes for across-sex analysis on FreeSurfer data using RF.

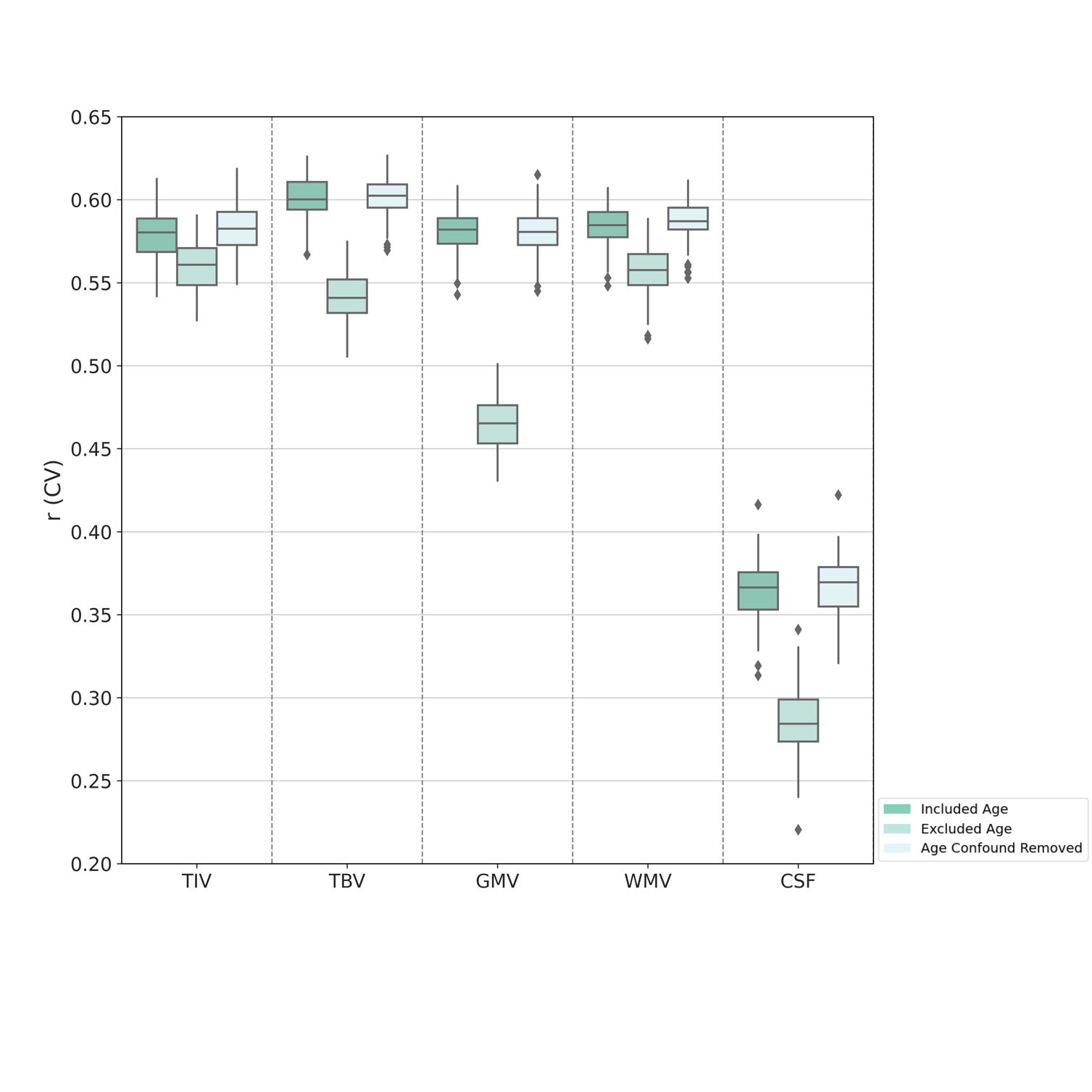

**Figure S12:** Impact of age on brain volumes for within-sex analysis on FreeSurfer data using linear SVM.

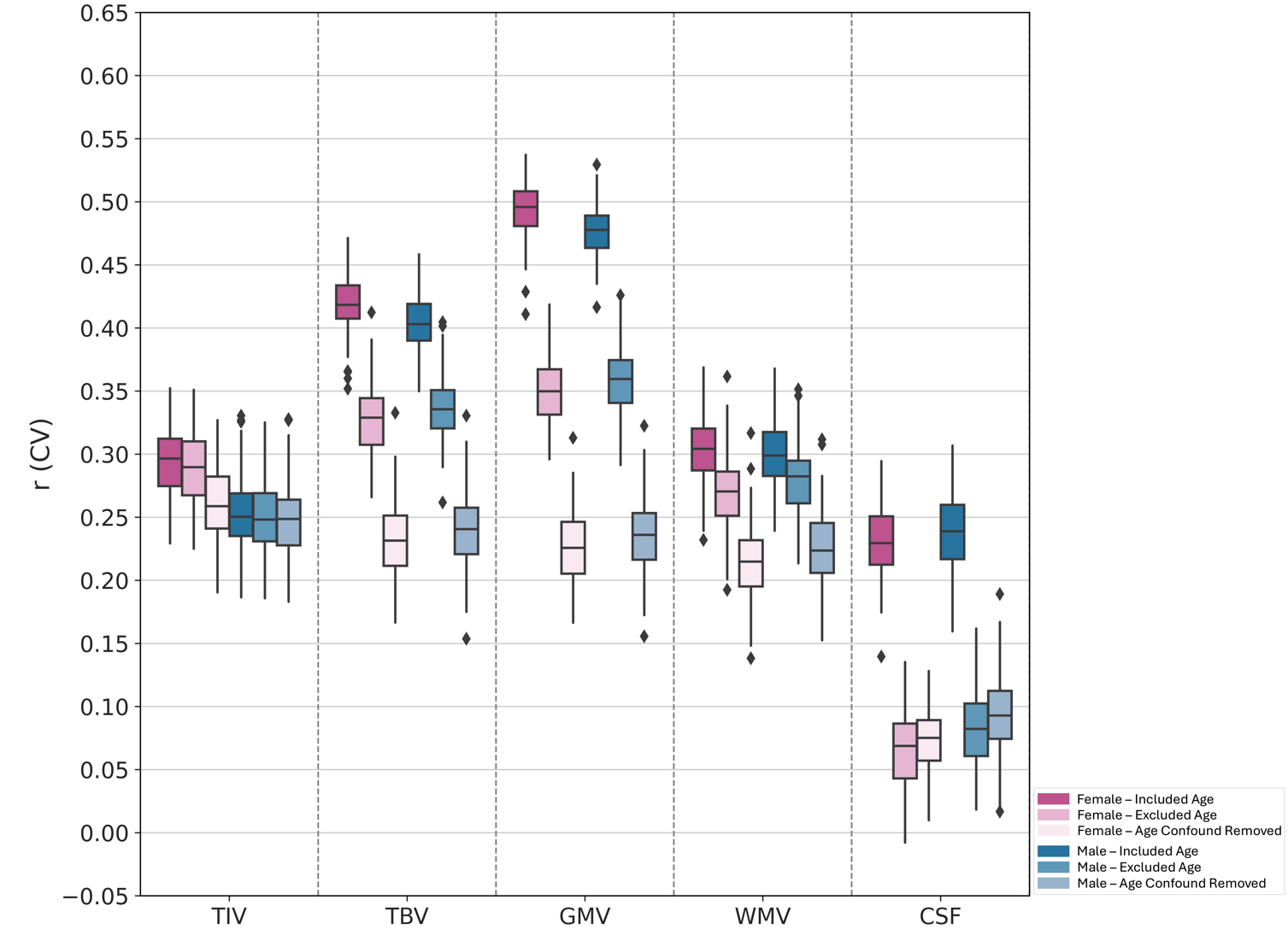

**Figure S13:** Impact of age on brain volumes for within-sex analysis on FreeSurfer data using RF.

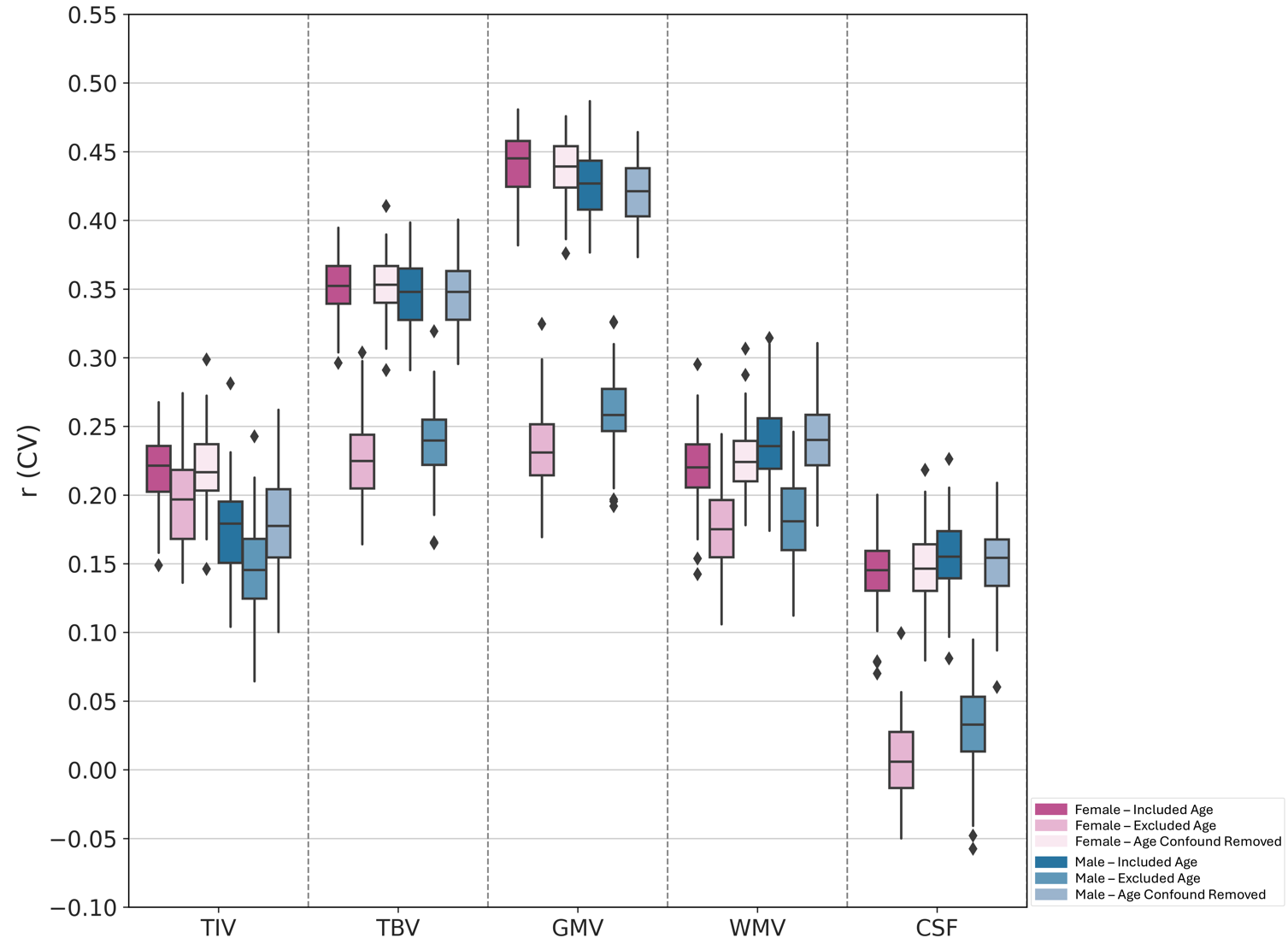
